# Encounter-state over-anchoring governs productive PETase binding on PET surfaces

**DOI:** 10.64898/2026.03.17.712535

**Authors:** Chengze Huo, Jun Wang, Xiakun Chu

## Abstract

Polyethylene terephthalate (PET) hydrolysis by *Ideonella sakaiensis* PETase (IsPETase) begins at a heterogeneous solid–liquid interface, yet the molecular basis of productive surface recognition remains poorly resolved. Here, we combined a Martini 3 coarse-grained PET model with GōMartini protein dynamics to investigate IsPETase binding to an extended PET surface. A four-state kinetic model, comprising unbound, encounter, docked, and pre-catalytic states, shows that productive binding is not limited by adsorption itself, but by a post-adsorption re-registration step that converts surface-bound encounter complexes into productively aligned configurations. The simulations reveal a stage-dependent role of conformational flexibility: flexible surface loops facilitate early capture, whereas excessive flexibility promotes misregistered hydrophobic contacts, over-stabilizes non-productive encounter states, and lowers the overall probability of productive commitment. Analysis of productive trajectories further identifies three microscopic reorientation modes by which the enzyme reaches the pre-catalytic state after adsorption. Comparative simulations of engineered PETase variants uncover a flexibility-driven speed–yield trade-off, in which increased flexibility accelerates successful binding events but reduces productive yield through encounter-state over-anchoring. Guided by this mechanism, we formulated a landscape-based design strategy that either weakens encounter-specific anchors or reinforces product-like contacts, leading to mutations that improve productive-binding yield. These results identify post-adsorption alignment as the key kinetic bottleneck in PETase surface recognition and provide a mechanistic framework for designing enzymes that operate at heterogeneous polymer interfaces.

## 1 Introduction

The rapidly growing accumulation of mismanaged plastic waste poses a persistent and escalating threat to ecosystems and to the long-term sustainability of the global biosphere [1–5]. Addressing this challenge requires not only reducing the production and release of virgin plastics, but also developing scalable recycling technologies capable of establishing circularity for dominant commodity polymers [6–10]. Among these materials, polyethylene terephthalate (PET) is particularly important because of its widespread use in packaging and textiles and its long-term persistence in the environment [4, 6, 8]. Although mechanical recycling is widely implemented, its effectiveness is often compromised by contamination, polymer-property deterioration, and downcycling. In parallel, many chemical recycling strategies remain energy-intensive and face substantial economic barriers to large-scale deployment [6–8, 11].

Enzymatic depolymerization has emerged as a promising strategy for PET recycling because, in principle, it enables closed-loop recovery by converting post-consumer PET into its constituent monomers under aqueous conditions and at comparatively mild temperatures, thereby facilitating re-polymerization into virgin-quality materials [6, 12–14]. A major advance in this field was the discovery of *Ideonella sakaiensis* and its two-enzyme PET degradation system, in which IsPETase initiates the hydrolysis of solid PET at the polymer surface to generate soluble intermediates, and MHETase subsequently converts these products into terephthalic acid (TPA) and ethylene glycol (EG) [15, 16]. Despite this promise, efficient PET depolymerization remains challenging because hydrolysis occurs at the interface of an insoluble, semi-crystalline substrate, where overall turnover depends not only on catalysis itself but also on productive adsorption, molecular orientation, substrate accessibility, and the dynamic conformational landscape of the enzyme-substrate complex [6, 12, 17–19]. In addition, effective depolymerization of real-world PET often benefits from operation near or above the glass-transition regime, where increased polymer-chain mobility enhances enzymatic accessibility but simultaneously imposes stringent demands on enzyme thermostability and operational robustness [6, 20–22]. Con-sequently, extensive engineering efforts, including rational design, computation-guided optimization, machine learning, and directed evolution, have been devoted to improving the stability and activity of PET hydrolases. Landmark variants such as ThermoPETase, DuraPETase, FAST-PETase, and Hot-PETase have substantially advanced PET depolymerization efficiency and temperature tolerance, bringing enzymatic recycling closer to industrial relevance [13, 20, 23–25].

At a fundamental level, PETase turnover can be viewed as three coupled stages: interfacial binding, hydrolytic cleavage, and product release [26]. While the chemical mechanism of ester-bond hydrolysis has been extensively characterized by QM/MM calculations and structural studies [27–29], the initial binding stage, which determines whether the enzyme can engage a recalcitrant solid substrate in a catalytically competent manner, remains far less well understood. Developing a predictive and mechanistic picture of how PET hydrolases achieve, or fail to achieve, productive binding on heterogeneous PET surfaces is therefore a central challenge for next-generation enzyme design [6, 12, 19, 30]. To date, most molecular-level insights have been derived from reductionist models based on soluble PET oligomers or single short polymer chains [31–34]. Although these studies have been valuable for resolving local binding interactions, they do not capture the heterogeneous adsorption landscape presented by realistic amorphous plastic surfaces. Moreover, simulations that explicitly include PET slabs often rely on enhanced-sampling schemes or supervised molecular dynamics [35, 36]. Such approaches are powerful for exploring thermodynamic basins, but they also impose external bias on the binding process, limiting their ability to resolve the authentic stochastic kinetics by which enzymes search, reorient, and commit to productive binding at a solid polymer interface [37].

To bridge this spatiotemporal gap, we established a computational platform based on the Martini 3 coarse-grained (CG) force field to simulate the complete binding process of IsPETase on a PET slab [38–44]. By analyzing a large ensemble of independent binding events across hundreds of microseconds of simulation, we reconstructed the full binding trajectory of IsPETase from bulk solution to productive engagement at the polymer interface. This analysis revealed three distinct mi-croscopic surface-reorientation modes that govern progression toward catalytically competent binding. The resulting pathway-resolved picture uncovers a stage-dependent role of conformational flexibility: dynamic surface loops facilitate initial capture, but excessive flexibility also promotes misregistered hydrophobic contacts that trap the enzyme in long-lived, non-productive encounter states. On the basis of this mechanism, we identify escape from these trapped states as the rate-limiting step for productive binding and formulate a rational design strategy that directly targets this kinetic bottleneck. By selectively destabilizing off-pathway interactions while reinforcing contacts unique to productive configura-tions, this framework led to the identification of three point mutations that significantly improve productive binding yield, thereby demonstrating the predictive power of a kinetics-informed design paradigm for enzymes operating at heterogeneous PET-liquid interfaces.

## 2 Results

### 2.1 CG modeling of PET using Martini 3

To balance computational efficiency with physical fidelity, we modeled polyethylene terephthalate (PET) chains using the Martini 3 CG force field (Figure 1A) [42, 43]. The CG representation of PET was constructed on the basis of the curated Martini small-molecule parameter database [43] and the fragment-based mapping guidelines of Martini 3 [42]. Following the Martini mapping philosophy, each PET repeat unit was partitioned into chemically distinct fragments, including the ester linkage (– C(– O) – O –), the aromatic ring, and the ethylene segment (– CH_2_ – CH_2_ –), and each fragment was assigned to an appropriate Martini 3 bead type according to its polarity, size, and chemical character. To enable automated construction of long polymer chains with Polyply [45], the PET topology was encoded as repeating CG building blocks, with terminal beads explicitly modified to account for end-group chemistry.

**Figure 1:**
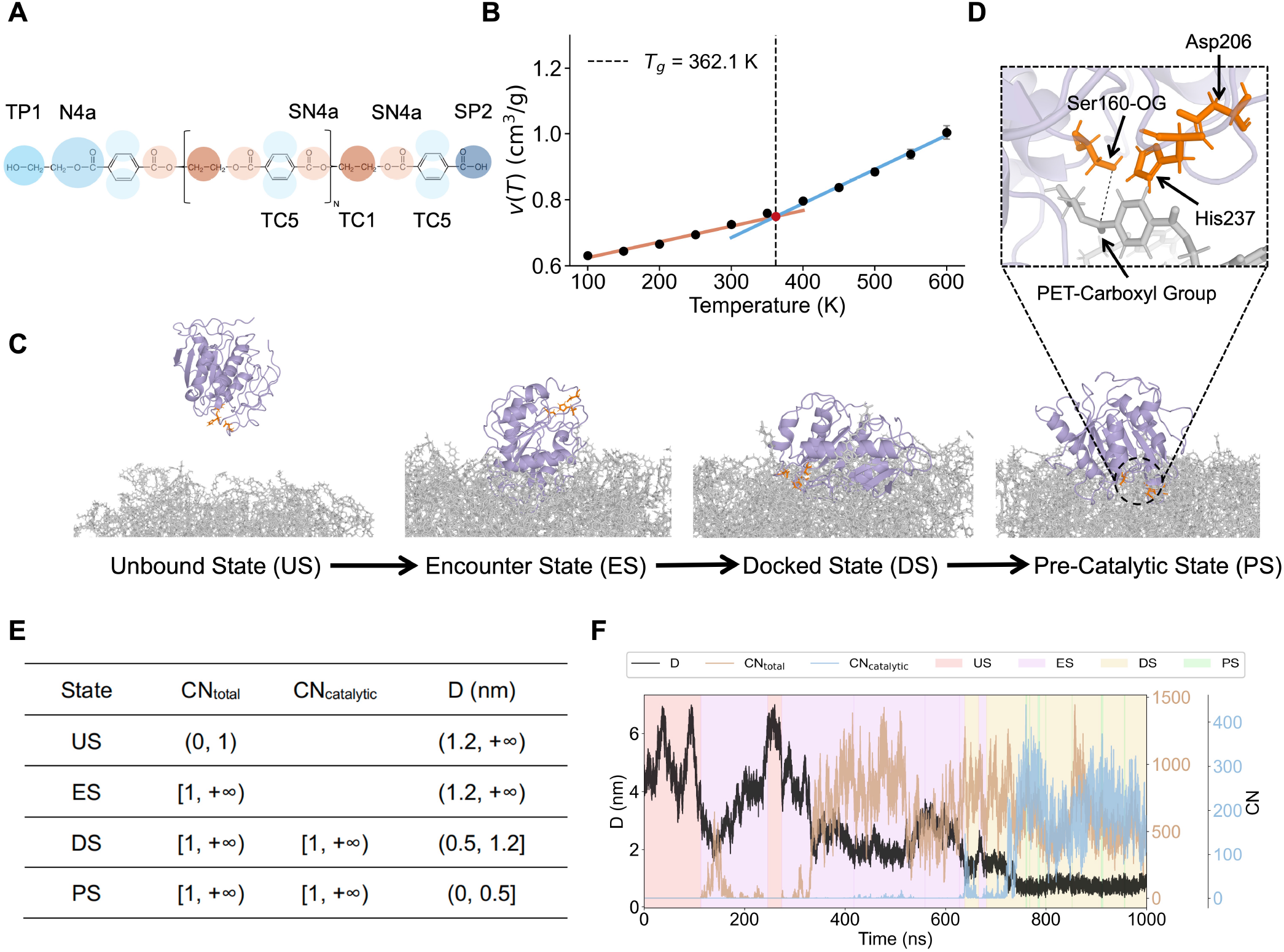
Construction of the computational platform and operational state definition for the binding of IsPETase to a PET slab. **(A)** Martini 3 coarse-grained (CG) mapping of a PET tetramer, showing the bead types assigned to the distinct chemical moieties. **(B)** Specific volume, *v*(*T*), of the PET slab as a function of temperature. Bilinear fits to the low- and high-temperature regimes give a glass-transition temperature of *T*_g_ = 362.1 K at their intersection (dashed lines). **(C)** Representative snapshots of the four binding states: unbound state (US), encounter state (ES), docked state (DS), and pre-catalytic state (PS). **(D)** Back-mapped atomistic view of the pre-catalytic state (PS), highlighting the catalytic triad (Ser160, His237, and Asp206) oriented toward a PET ester carbonyl group. **(E)** Criteria used for state classification based on three metrics: the total protein-PET contact number, CN_total_; the catalytic-pocket contact number, CN_catalytic_; and the active-site proximity, *D*. Here, *D* is defined as the Euclidean distance between the side-chain bead (SC1) of Ser160 and the centroid of the five nearest PET beads. **(F)** Representative trajectory showing *D*(*t*) (black), CN_total_(*t*) (orange), and CN_catalytic_(*t*) (blue). Background shading indicates the assigned binding state over time, illustrating transitions driven by the coupled formation of protein-surface contacts and reduction of the active-site distance.

Within this representation, nonbonded interactions were determined by the assigned Martini bead types, whereas bonded and torsional potentials were calibrated against all-atom simulations of PET dimers (see details in Methods). After iterative refinement, the resulting CG parameter set reproduced the structural distributions of the atomistic reference with good accuracy (Figures S1 and S2). In particular, the average solvent-accessible surface area (SASA) differed from the all-atom model by only 1.23% (Figure S2), indicating close agreement between the two levels of representation. Using this validated Martini 3 model, we next constructed a 10 × 10 × 5 nm^3^ PET slab with Polyply [45], comprising 200 PET chains with 10 monomer units per chain, and performed molecular dynamics simulations to estimate its glass-transition temperature (*T*_g_). The resulting *T*_g_ of 362.1 K (Figure 1B) agrees well with the experimental value of approximately 350 K [46–48], supporting that the developed Martini 3 model captures the key structural and thermophysical properties of PET and is suitable for investigating the binding of PETase to a PET slab.

### 2.2 A four-state kinetic framework for IsPETase binding to PET

To allow IsPETase to sample its native conformational ensemble during surface recognition, we modeled the enzyme using the GōMartini 3 force field [40, 44], which preserves the native fold while permitting large-scale collective motions. Building on the calibrated Martini 3 parameters for both PET and IsPETase, we established an explicit protein-surface simulation platform to resolve adsorption and binding on an extended PET substrate. All subsequent protein-PET simulations employed the same PET slab described above. The wild-type IsPETase structure (PDB: 5XJH) [33] was initially placed at least 4 nm away from the PET surface (Figure S3), well beyond the range of direct interactions, thereby allowing spontaneous approach and binding from bulk solution without steering during the simulations. We then carried out ∼100 independent simulations to generate an ensemble of binding trajectories with robust statistical coverage (Table S2).

To quantitatively characterize the binding dynamics, we defined a minimal set of reaction coordinates that jointly describe (i) global adsorption, (ii) active-site engagement, and (iii) local proximity of the catalytic nucleophile to the polymer surface. Specifically, we monitored the total protein-PET contact number, CN_total_, and the catalytic-pocket contact number, CN_catalytic_, both defined using a distance-based contact criterion (see details in Methods). We further introduced an active-site proximity metric, *D*, defined as the Euclidean distance between the side-chain bead (SC1) of Ser160 and the centroid of the five nearest PET beads. The catalytic pocket was represented by ten residues that define the binding cleft and catalytic region, including Tyr87, Trp159, Ser160, Met161, Trp185, Asp206, Ile208, His237, Trp238, and Asn241 (Figure S3B) [33]. On the basis of the joint evolution of CN_total_, CN_catalytic_, and *D*, we partitioned the binding process into four operational states (Figure 1C-F), enabling consistent state assignment across the full trajectory ensemble.

This decomposition reveals a multi-step binding pathway (Figure 1C-F). In the Unbound State (US), the enzyme diffuses in bulk solution without persistent contacts with the substrate. Initial nonspecific adsorption gives rise to the Encounter State (ES), in which CN_total_ increases while the catalytic pocket remains largely disengaged from the surface. Although adsorbed, the enzyme in ES is generally not catalytically relevant, because its orientation is sub-optimal and the active-site groove does not necessarily face the polymer. The system subsequently enters the Docked State (DS), where surface diffusion and reorientation allow the catalytic pocket to establish sustained contacts with the PET surface. Finally, trajectories reach the Pre-Catalytic State (PS), defined by close active-site proximity together with catalytic-pocket engagement. In PS, the catalytic triad (Ser160, His237, and Asp206) is oriented toward a PET ester carbonyl group in a geometry consistent with a catalytically competent configuration (Figure 1D). The corresponding time series (Figure 1F) further shows that transitions between these states are governed by the coupled formation of protein-surface contacts and the progressive reduction of *D*.

### 2.3 Conformational dynamics drive stage-dependent surface recognition

We next examined how conformational dynamics shape the multi-step binding pathway of IsPETase. State-resolved root-mean-square fluctuation (RMSF) analysis reveals a monotonic decrease in global flexibility from the US to the PS (Figure 2A-C), indicating progressive restriction of conformational degrees of freedom as the enzyme engages the surface and approaches a catalytically competent configuration (Figure 2D-F).

**Figure 2:**
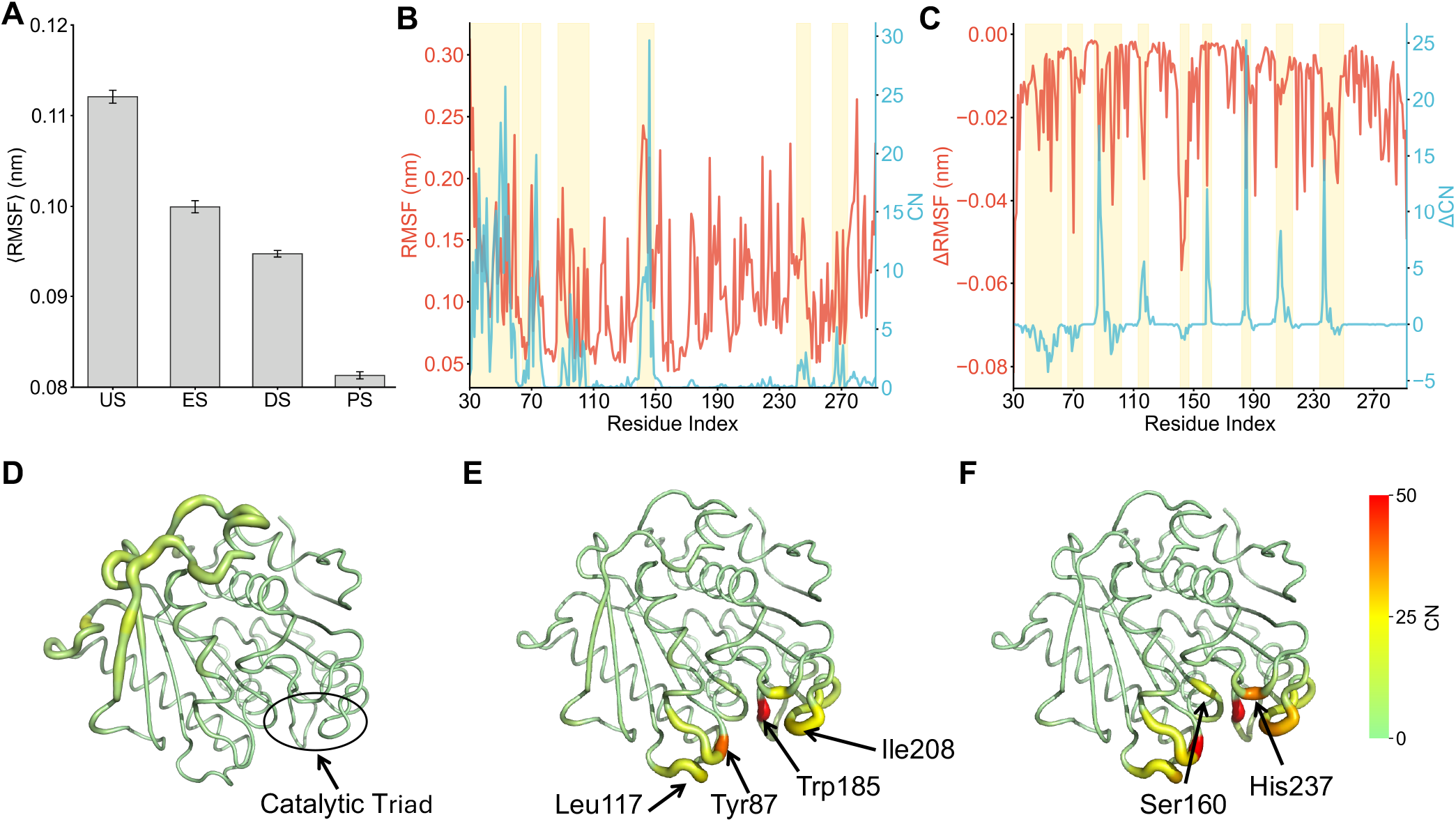
Stage-dependent role of conformational flexibility and evolution of interfacial contacts during IsPETase binding. **(A)** Average protein RMSF decreasing monotonically from the US to the PS. This indicates progressive rigidification along the binding pathway. **(B)** Fly-casting-like capture during the encounter stage. Residue-wise RMSF in the US (red) and contact frequency in the ES (blue) are plotted together for comparison. Shaded regions mark major overlap peaks, showing that highly flexible segments in solution preferentially mediate initial surface anchoring. **(C)** Residue-wise reorganization of flexibility and interfacial contacts during binding maturation from the DS to the PS. Changes in flexibility (ΔRMSF = RMSF_PS_ − RMSF_DS_, red) and contact number (ΔCN = CN_PS_ − CN_DS_, blue) reveal a coordinated redistribution of dynamics and protein-surface interactions as the enzyme approaches a catalytically competent configuration. **(D-F)** Residue-wise contact mapping between IsPETase and the PET slab in the ES, DS, and PS, respectively. The protein is shown as a tube, with both thickness and color (green to red) scaled by the mean contact number. Labeled residues highlight the spatial shift of interaction hotspots from early capture regions toward the catalytic vicinity as binding progresses.

During the early stage of recognition, local flexibility appears to facilitate substrate capture. In the ES, the residue-level contact profile is enriched in loop-dominated regions (Figure 2B,D and Figure S4), indicating that structurally mobile segments contribute disproportionately to initial adsorption. Consistent with a fly-casting capture mechanism [49, 50], comparison of the intrinsic flexibility in the US (RMSF, red) with the ES contact frequency (blue) shows that the most flexible segments in solution coincide with the dominant contact hotspots upon encounter (highlighted regions in Figure 2B). These flexible loops therefore appear to enlarge the effective capture radius, enabling transient surface anchoring before productive orientation is established.

In contrast, the transition from the DS to PS is characterized not by uniform quenching of dynamics, but by a functional redistribution of flexibility and interfacial contacts. Structural contact analysis (Figure 2C, 2E, 2F and Figure S4) identifies a subset of hydrophobic and aromatic residues, most prominently Tyr87, Leu117, Trp185, and Ile208, that progressively accumulate PET contacts and act as surface anchors to stabilize the docking orientation. At the same time, residues in the catalytic core, including Ser160 and His237, also gain contacts while becoming less flexible, consistent with the local rigidification required to establish a pre-catalytic geometry.

Importantly, contact-induced rigidification is not uniformly coupled to increased surface interaction across the entire protein. As shown in Figure 2C, several regions that exhibit pronounced RMSF reduction, including the terminal segments and loop regions around Tyr146, simultaneously lose PET contacts during the DS→PS transition. These segments appear to function primarily in exploratory recogni-tion during the US→ES stages and then become progressively disengaged as binding proceeds. Productive binding is therefore consolidated by a more localized anchoring network, while early-recognition segments relinquish transient adsorption roles and undergo passive conformational quenching. Together, these results support a stage-dependent and spatially heterogeneous role of conformational dynamics: flexibility promotes long-range surface capture and early adsorption, whereas selective rigidification stabilizes anchoring and catalytic residues required for productive engagement.

To place these observations in a global conformational context, we performed principal component analysis (PCA) on the protein trajectories and constructed state-resolved free-energy landscapes (Figure S5). The US ensemble spans multiple basins, indicating a diverse population of pre-existing conformers (Figure S5A). Upon surface engagement, this distribution progressively narrows, and the bound ensembles become increasingly enriched in a subset of basins that dominate the DS and PS states (Figure S5B-E). Notably, the basins most strongly populated in PS are already accessible in the US ensemble, indicating that productive binding contains a clear conformational-selection component rather than arising solely from binding-induced remodeling [51]. At the same time, the progressive reduction in flexibility upon binding indicates additional stage-specific stabilization after surface engagement. This population shift suggests that productive binding may be enhanced by biasing the unbound ensemble toward PS-like conformations while disfavoring conformers that preferentially enter non-productive encounter states.

### 2.4 Kinetic partitioning reveals the thermodynamic origin of trap states

Not all surface-bound trajectories proceed to a catalytically competent configuration. Across the independent trajectories, surface engagement is highly probable, with 96.08% of trajectories reaching the ES and 85.29% further progressing to the DS, whereas only 56.86% ultimately reach the PS. This substantial loss in PS yield indicates that binding is kinetically partitioned into a productive pathway and a competing non-productive pathway (inset, Figure 3A). This behavior is consistent with previous simulation studies suggesting that IsPETase often adsorbs to PET in non-productive orientations and that catalytically competent poses are only weakly favored energetically [36].

**Figure 3:**
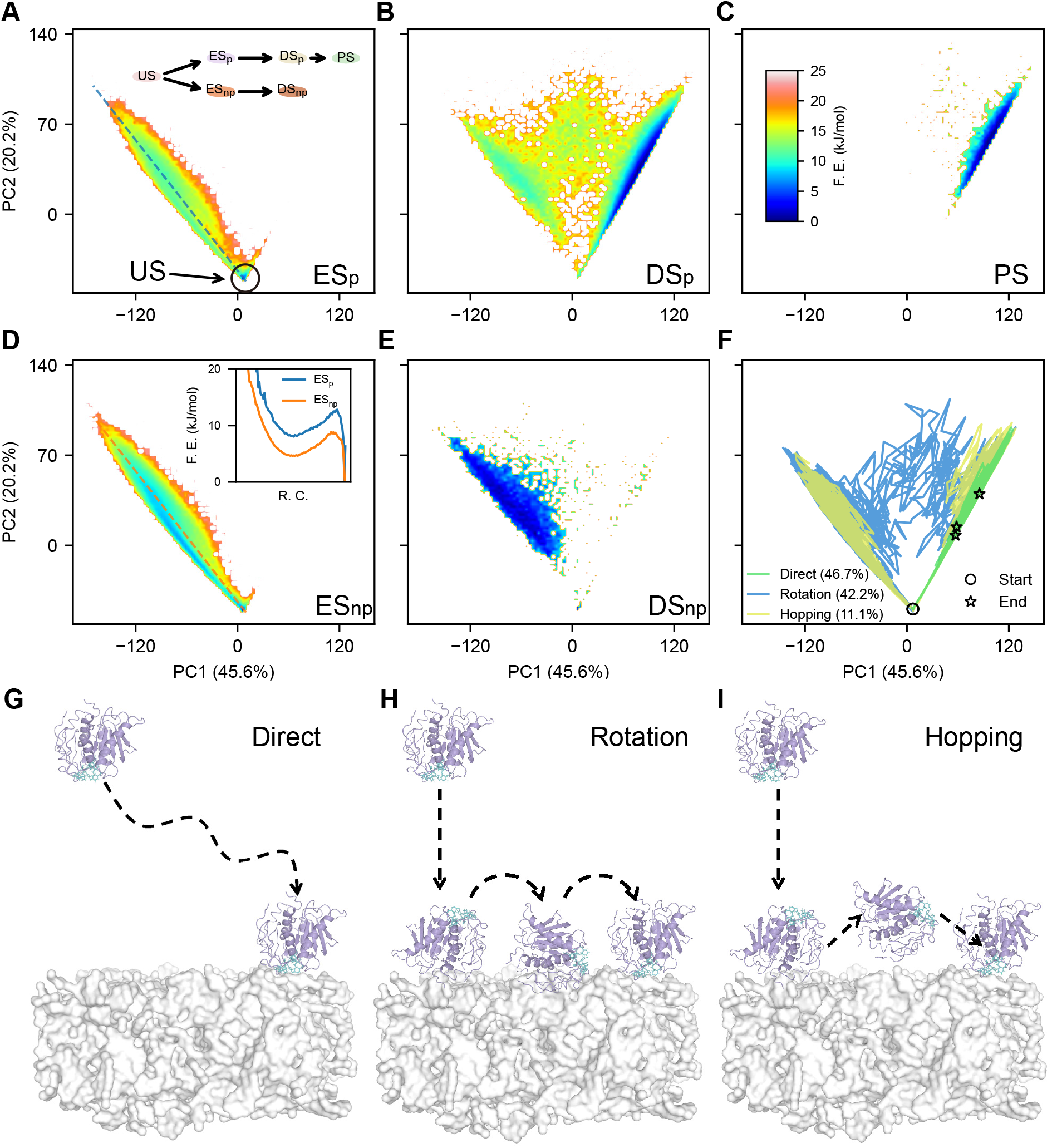
Free-energy landscapes and microscopic pathways underlying productive and trapped Is-PETase binding on PET. **(A–C)** Free-energy landscapes (FELs) in the space of the first two principal components (PC1 and PC2), obtained from principal component analysis of residue-wise contact distributions, for the productive ensembles ES_p_, DS_p_, and PS. Along the productive route, the system evolves from broad ES_p_ and DS_p_ basins toward the distinct contact pattern of PS. The inset in **(A)** schematically summarizes the bifurcated binding mechanism, consisting of a productive pathway (US→ES_p_→DS_p_→PS) and a competing trapped pathway (US→ES_np_→DS_np_), in which misoriented contact patterns stabilize non-productive ensembles on the simulated timescales. **(D**,**E)** FELs of the non-productive ensembles ES_np_ and DS_np_. The inset in **(D)** shows one-dimensional free-energy profiles projected along the transition direction (dashed line; reaction coordinate, R.C.), illustrating that ES_np_ occupies a deeper local minimum than ES_p_, consistent with kinetic trapping. **(F)** Productive trajectories projected onto the same PCA space reveal three microscopic conversion modes from ES_p_ to PS: Direct, Rotation, and Hopping. **(G−I)** Representative schematics and observed frequencies of the three productive modes: **(G)** Direct (46.7%), **(H)** Rotation (42.2%), and **(I)** Hopping (11.1%).

Residue-resolved contact analysis distinguishes these two pathways at the structural level. Trajectories that reach PS undergo pronounced reorganization of interfacial contacts during the ES→DS transition, whereas trajectories that fail to reach PS retain DS contact patterns that remain highly similar to those of ES, indicating limited progression beyond the initial encounter configuration (Figures S6 and S7). On this basis, we define a non-productive encounter ensemble, ES_np_, that can further evolve into a misdocked state, DS_np_, but remains kinetically disconnected from the productive route on the simulated timescales (inset, Figure 3A). Consistent with this interpretation, productive trajectories pass relatively quickly through ES_p_ and DS_p_, whereas non-productive trajectories exhibit prolonged residence in the encounter-like ES_np_ regime, reflecting frustrated progression toward productive docking and active-site engagement (Figures S8A,B).

To rationalize this kinetic partition thermodynamically, we constructed free-energy landscapes by performing principal component analysis on residue-wise contact distributions (Figure 3). Along the productive route (Figure 3A–C), the landscape reveals a discrete reorganization of contact patterns rather than a smooth, continuous convergence. ES_p_ and PS occupy topologically separated regions in contact-PCA space, and the transition between them occurs primarily during the ES_p_→DS_p_ stage. DS_p_ spans an intermediate region that bridges these basins, consistent with its role as a surface-reorientation and contact-remodeling state.

By contrast, ES_np_ and DS_np_ (Figure 3D,E) remain confined to a region of contact space that is disconnected from the productive PS basin. Importantly, the one-dimensional free-energy profile projected along the transition direction (Figure 3D, inset) shows that ES_np_ occupies a deeper local minimum than ES_p_, indicating that non-productive binding is stabilized by a thermodynamic sink formed by strong-but-wrong contacts. The low PS yield therefore does not arise from insufficient surface affinity, but from the rarity of trajectories that successfully reorganize their interfacial contacts after adsorption and cross the ES→DS bottleneck required to access the PS basin.

Among the trajectories that do reach the PS, we identify three distinct microscopic modes for the ES_p_→PS conversion (Figure 3F−I). In the “Direct” mode, the enzyme lands in a near-productive orientation upon first stable adsorption and reaches the PS with minimal subsequent rearrangement. In the “Rotation” mode, the enzyme remains surface-bound while undergoing lateral diffusion and gradual reorientation, allowing productive alignment to emerge without detachment. In the “Hopping” mode, transient desorption and re-adsorption reset the interfacial registry, providing an escape route from misoriented contact patterns. The coexistence of these three modes shows that productive binding is governed not by a single dominant route, but by the ability of the enzyme to reorganize interfacial contacts after adsorption. In this sense, successful trajectories are distinguished primarily by their capacity to escape misoriented encounter states through surface reorientation or registry resetting, highlighting post-adsorption reorganization as the key determinant of the PS formation.

### 2.5 Encounter-state over-anchoring creates a flexibility-driven speed–yield trade-off

To test whether the binding mechanism identified for the wild type (WT) generalizes across sequence space, and to quantify how conformational flexibility modulates productive binding, we simulated three engineered IsPETase variants with distinct flexibility profiles (Thermo, FAST, and Hot) using the same simulation protocol and state definitions applied to WT. For each variant, we collected a large ensemble of independent trajectories (Table S2), enabling direct comparison of binding kinetics and state populations across systems.

All variants readily formed encounter complexes, with fractions of trajectories reaching the ES comparable to that of WT (Figure 4A,B). Their overall productive yield, however, was reduced: fewer trajectories progressed to the DS and ultimately to the PS. We traced this loss primarily to the early adsorption regime. Across the variants, the mean contact number within the ES ensemble is inversely correlated with the probability of reaching PS (Figure 4C), indicating that excessive nonspecific contacts formed immediately after landing suppress productive binding. Consistent with this interpretation, the more flexible variants exhibit prolonged residence in the ES (Figure S8A,B), suggesting that over-anchoring kinetically traps misoriented encounter conformations and reduces the likelihood of subsequent progression toward the PS.

**Figure 4:**
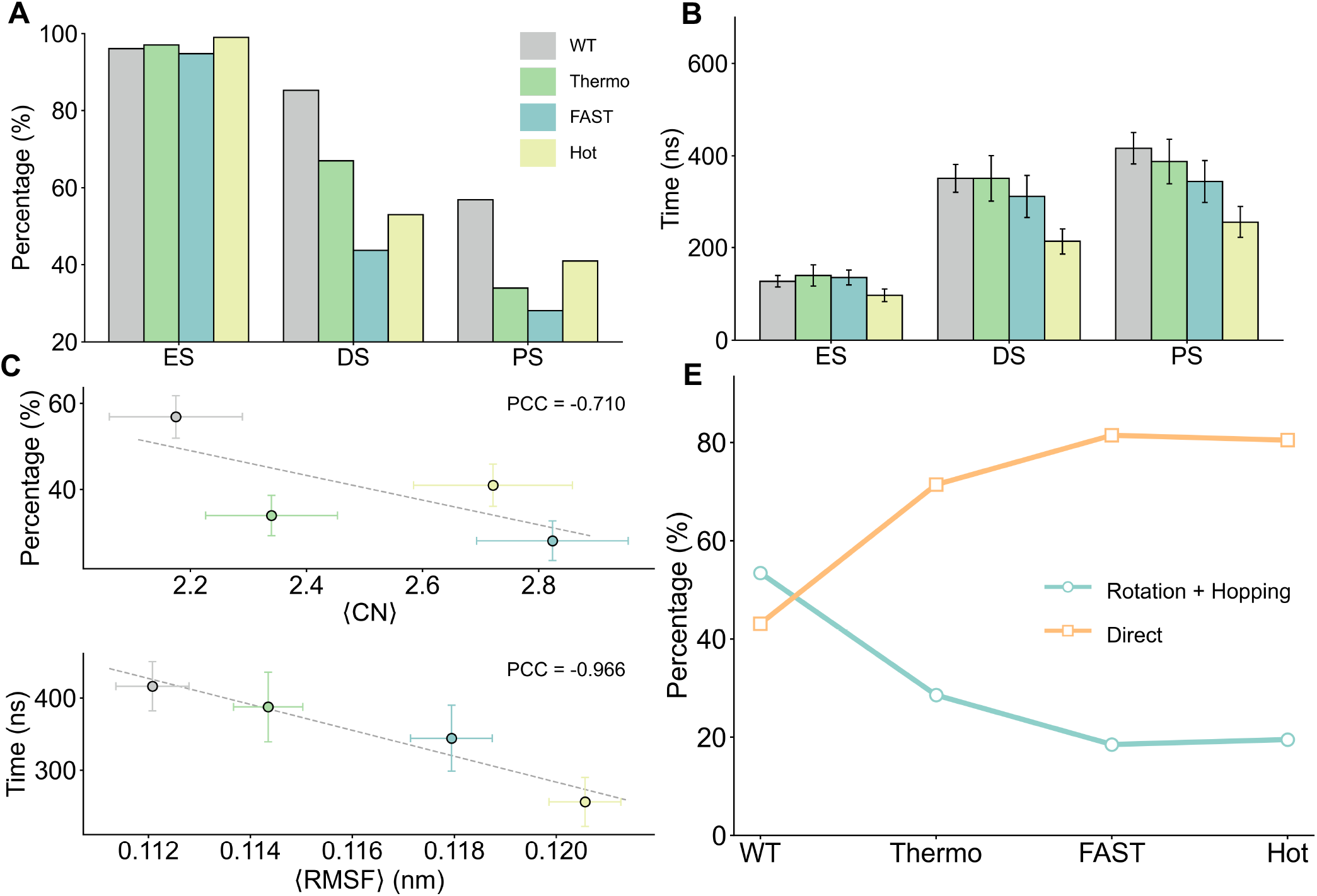
Encounter-state over-anchoring underlies a flexibility-driven speed-yield trade-off across PETase variants. **(A)** Fractions of trajectories reaching the ES, DS, and PS for WT and three engineered variants (Thermo, FAST, and Hot), calculated from independent trajectory ensembles for each system. **(B)** Mean first-arrival times to the ES, DS, and PS, computed over trajectories that successfully reach the corresponding state. **(C)** Relationship between the mean contact number in the ES ensemble and the overall probability of reaching PS across variants. The dashed line shows a linear fit; PCC denotes the Pearson correlation coefficient. **(D)** Relationship between unbound-state flexibility, measured by the average RMSF in the US ensemble, and the mean first-arrival time to PS, showing that higher initial flexibility is associated with faster productive binding among successful trajectories. **(E)** Composition of productive binding modes among PS-reaching trajectories. With increasing flexibility, the fraction of trajectories that reach PS through reorientation-dependent pathways (Rotation + Hopping) decreases, whereas near-direct landing events (Direct) become increasingly dominant, consistent with faster but lower-yield productive binding in the more flexible variants.

At the same time, the flexible variants reach the PS more rapidly than WT among the subset of trajectories that do succeed (Figure 4B). Specifically, the mean first-arrival time to the PS decreases as the flexibility of the unbound-state ensemble increases, yielding a negative trend across the four tested variants between the US RMSF and the PS first-arrival time (Figure 4D). This apparent acceleration does not contradict the lower overall yield; rather, it reflects a shift in which microscopic routes remain accessible once early adsorption becomes stronger.

Pathway decomposition of the PS-reaching trajectories reveals a systematic redistribution of productive modes with increasing flexibility. As flexibility increases, the fraction of trajectories that require substantial post-adsorption reorientation, namely Rotation and Hopping, decreases, whereas near-direct landing becomes progressively dominant (Figure 4E). Because Direct trajectories bypass extended interfacial adjustment, they reach the PS more rapidly than routes that depend on reorientation after adsorption (Figure S8C,D), thereby explaining the shorter time-to-PS among the successful subset. These results suggest that higher flexibility promotes richer and more persistent ES contact patterns (Figures S6 and S7) that become increasingly difficult to reorganize once established. As a consequence, productive binding becomes progressively more dependent on landing in a near-productive orientation from the outset, rather than on reorientation-enabled rescue after adsorption.

Taken together, these results establish a flexibility-driven speed–yield trade-off governed by encounter-state over-anchoring. Increased flexibility can accelerate productive binding when a near-productive landing occurs, but it simultaneously strengthens nonspecific adsorption in the ES, prolongs trapping in encounter-like configurations, and suppresses reorientation-dependent rescue pathways, thereby lowering the overall PS yield. Efficient productive binding therefore requires a balance between conformational mobility and controlled surface anchoring, directly motivating the design strategy developed below.

### 2.6 Rational design targeting the rate-limiting step

Our mechanistic analysis shows that productive binding of IsPETase to PET is not limited by initial surface encounter formation, but by the subsequent re-registration step that converts the ES into a productively aligned bound configuration capable of progressing through the DS toward the PS. Although adsorption occurs readily, a substantial fraction of trajectories become trapped in misregistered encounter complexes, where non-native contacts stabilize deep kinetic basins and suppress the rotational or hopping rearrangements required for productive docking. The ES→DS transition therefore emerges as the dominant rate-limiting step in the binding pathway.

Guided by this picture, we considered two complementary strategies to relieve the ES→DS bottleneck (Figure 5A). The first strategy seeks to stabilize product-like configurations by selectively reinforcing contacts that persist in DS and PS, thereby deepening the product basin and lowering the ES–DS free-energy barrier. The second strategy targets the opposite limit by weakening encounter-specific interactions that stabilize misregistered complexes and generate kinetic traps. By reducing the depth of the encounter basin, this approach increases the probability that surface-adsorbed enzymes can escape non-productive registrations and undergo the rearrangements required to reach the DS.

**Figure 5:**
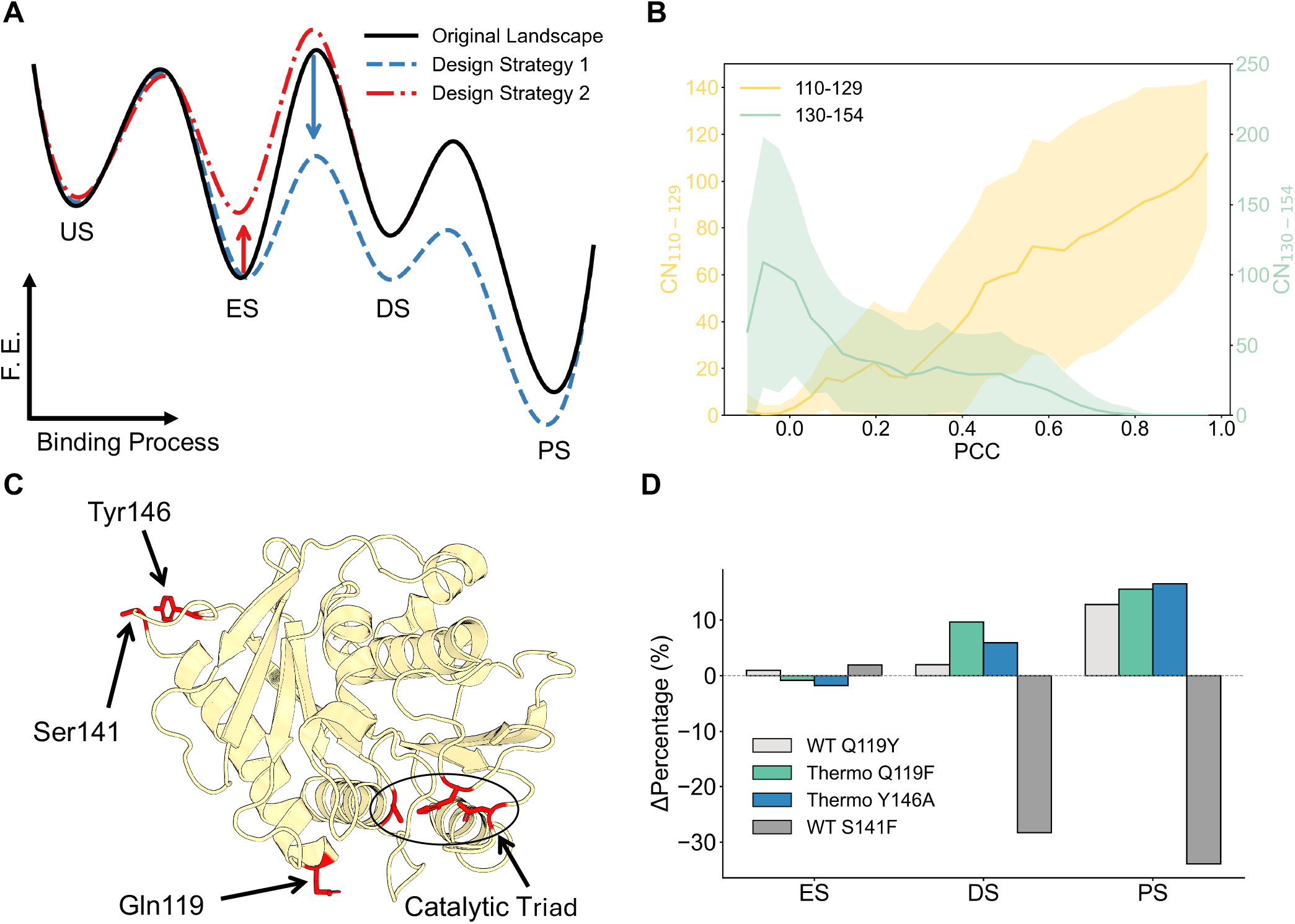
Domain-resolved contact reorganization identifies residue-level targets for rational design of the ES→DS transition. **(A)** Schematic free-energy profiles along the binding pathway from the US to the PS. The original landscape (black) contains an ES basin and an ES-DS barrier that limits productive docking. **Design strategy 1** (blue dashed line) stabilizes the DS and PS by strengthening product-like contacts, thereby lowering the ES-DS barrier and deepening the DS and PS basin. **Design strategy 2** (red dash-dot line) destabilizes the ES by weakening encounter-specific anchors that create kinetic traps, thereby biasing trajectories toward the DS. Arrows indicate the intended direction of landscape modulation rather than quantitative free-energy changes. **(B)** PCC-dependent mean contact sums for two loop regions (residues 110-129 and 130-154). For each PCC bin, where PCC is the Pearson correlation coefficient between the instantaneous residue-PET contact profile and the DS reference, the solid line shows the binned mean contact sum and the shaded band denotes the within-bin standard deviation. Separate *y*-axes are used for the two residue ranges. **(C)** Structural mapping of the three design residues, Gln119, Ser141, and Tyr146 (red), on IsPETase, shown relative to the catalytic triad. **(D)** State-resolved changes induced by the designed substitutions, reported as absolute differences (percentage points) in the fractions of trajectories reaching the ES, DS, and PS relative to the corresponding parent background (WT or Thermo; see details in Methods). Variants that strengthen product-like contacts (Q119Y/Q119F) or weaken encounter-specific anchoring (Y146A) shift outcomes toward DS and PS, whereas increasing encounter propensity (S141F) illustrates the capture-trap trade-off.

To localize these strategies at the residue level, we analyzed how residue–PET contact patterns reorganize along the ES→DS transition using a reaction coordinate defined by the similarity between instantaneous contacts and the DS reference ensemble (Figure S9). As trajectories progress toward productive docking, this coordinate increases monotonically, allowing us to distinguish surface regions that engage early and remain populated during re-registration from those that contribute primarily to ES. This analysis reveals two surface segments with contrasting functional roles (Figure 5B). Contacts involving the 110−129 loop emerge early during the ES→DS transition and remain strongly populated in DS and PS, indicating that this region directly stabilizes product-like surface registrations. By contrast, the 130−154 loop forms substantial contacts in ES that decay sharply as trajectories approach DS, suggesting that interactions mediated by this segment are characteristic of misregistered encounter complexes and act as kinetic anchors that hinder productive reorientation.

We first designed an explicit negative control to test whether accelerating surface capture alone improves productive binding. On the basis of encounter-state contact signatures (Figure S4A), we introduced a hydrophobic/aromatic substitution at the encounter interface (WT-S141F) to enhance nonspecific surface affinity (Figure 5C). As expected, this mutation shortened the unbound search phase and increased the fraction of trajectories reaching ES. However, this apparent gain proved counterproductive: strengthened encounter contacts deepened misregistered basins and reduced the fraction of trajectories progressing to DS and PS by an absolute 30 percentage points (Figure 5D). This result shows that once encounter formation is already efficient, further strengthening nonspecific adsorption is not a useful design objective.

We next evaluated design strategy 1 by selectively reinforcing product-like contacts. Within the 110–129 loop, Gln119 emerged as a leverage point: although the loop as a whole engages PET strongly, this position underutilizes hydrophobic and aromatic packing against the polymer surface. We therefore introduced an aromatic substitution (Q119Y) in the WT background to stabilize contacts characteristic of the DS and PS. This mutation increased the PS yield by an absolute 12.8 percentage points relative to WT, while leaving the mean first-arrival time among successful trajectories essentially unchanged (Figure S10), indicating that the mutation improves the probability of productive docking without imposing an additional kinetic penalty. Applying the same design logic to the more flexible Thermo variant, Thermo-Q119F partially rescued its pronounced ES→DS attrition, increasing DS and PS yields by absolute 9.6 and 15.6 percentage points, respectively.

Finally, we tested design strategy 2 by weakening an encounter-specific anchor within the 130–154 loop. Residue-level variance analysis comparing PS-reaching and non-PS trajectories identified Tyr146 as a dominant contributor to encounter-state stabilization (Figure S4D), consistent with its aromatic side chain acting as a strong but misdirecting surface anchor (Figure 5C). Removing this interaction (Thermo-Y146A) produced the largest improvement among all designs, increasing PS yield by an absolute 16.49 percentage points (Figure 5D). Within the present simulation framework, these results support weakening encounter-specific anchoring as an effective route to improving productive binding.

Taken together, these results establish a coherent and experimentally testable framework for landscape-guided enzyme surface adaptation. Simply increasing encounter propensity accelerates capture but aggravates kinetic trapping, whereas selectively strengthening product-like contacts or weakening encounter-specific anchors enhances productive binding while preserving the intrinsic timescale of successful ES→PS progression.

## 3 Discussion and Conclusions

In this work, we establish a scalable simulation framework for resolving enzyme recognition at a solid–liquid polymer interface by combining Martini 3 with a GōMartini-based description of protein dynamics. This framework enables long-timescale, high-statistics sampling of IsPETase binding on an extended PET slab and supports a compact four-state kinetic model (US→ES→DS→PS) that separates initial surface capture from post-adsorption alignment and pre-catalytic engagement. More importantly, it shows that productive PET recognition is governed not by adsorption alone, but by the ability of the enzyme to escape misregistered encounter complexes and progress through the ES→DS bottleneck toward the PS.

This result sharpens the current picture of interfacial PET hydrolysis. Because PET is an insoluble substrate that can only be accessed at the interface, overall enzymatic performance depends not only on the chemistry of ester-bond cleavage, but also on whether the enzyme can form a productive surface registration after landing [26, 52]. Recent studies have emphasized the prevalence of non-specific binding populations and the limited energetic bias toward catalytically relevant poses on PET-like surfaces [35, 36, 52–54]. Our simulations provide a molecularly resolved explanation for this behavior by showing that a large fraction of trajectories become trapped after adsorption in misregistered encounter states stabilized by strong-but-wrong contacts. Productive binding therefore depends on a post-adsorption search problem: the enzyme must reorganize its interfacial registry so that the binding cleft becomes aligned with locally accessible PET segments. In this sense, the ES→DS transition is not simply an intermediate refinement step, but the dominant kinetic gate that controls commitment to catalysis.

The pathway analysis further shows that this commitment process is microscopically heterogeneous. We resolve three distinct productive routes: near-productive landing, in-place rotation while remaining adsorbed, and hopping through transient desorption and re-adsorption. This result extends recent work on PET hydrolase binding/entry pathways [35] by showing that productive recognition at an extended polymer interface cannot be described as a single downhill adsorption event. Instead, it is governed by the ability of the enzyme to either preserve a favorable initial registry or recover from an unfavorable one through local surface reorganization. The thermodynamic analysis reinforces this interpretation: non-productive encounter ensembles occupy deeper local minima than their productive counterparts, indicating that low productive yield arises not from insufficient affinity, but from over-stabilization of off-pathway states. This state-specific view also refines the Sabatier-like interpretation of polyester hydrolysis [26]: the relevant design variable is not binding strength in the abstract, but how binding energy is distributed across the ES, DS, and PS.

A second major implication of our results is that conformational flexibility plays a two-edged, stage-dependent role in interfacial recognition. During the US→ES stage, flexible loops facilitate early capture, consistent with general recognition frameworks in which conformational breadth accelerates target search and selection. Once the enzyme is adsorbed, however, the same flexibility can become counterproductive: it promotes richer and more persistent short-range contact patterns, prolongs residence in misregistered ES configurations, and suppresses the rotational or hopping rearrangements required to escape them. This mechanism explains the flexibility-driven speed-yield trade-off observed across the engineered variants. More flexible variants reach PS faster among successful trajectories, yet they exhibit lower overall productive yield because productive binding becomes increasingly dependent on near-direct landing rather than on reorientation-enabled rescue. Rather than supporting a monotonic “more flexibility → better function” narrative, our data indicate that productive interfacial binding requires a balance between conformational mobility and controlled anchoring. The relevant design goal is therefore not global flexibilization, but redistribution of flexibility: preserve search-enabling dynamics while constraining the residues and loops that most strongly stabilize off-pathway encounter states [36, 55].

These mechanistic insights naturally reframe enzyme optimization as a landscape-engineering problem. The key design objective is to increase the commitment probability across the rate-limiting ES→DS bottleneck and thereby raise the fraction of trajectories that ultimately reach the PS. Within this framework, the two strategies tested here are mechanistically complementary. Strengthening product-like contacts deepens the DS/PS basin and stabilizes productive surface registrations, whereas weakening encounter-specific anchors reduces the depth of the ES trap and facilitates escape toward the DS. The designed mutants support both principles. The negative-control mutation that enhances encounter propensity improves initial capture but markedly decreases productive yield, confirming that stronger nonspecific adsorption alone is not a useful design objective once encounter formation is already efficient. By contrast, mutations that reinforce DS/PS-like contacts or weaken encounter-only anchors increase productive binding without imposing an obvi-ous kinetic penalty on successful trajectories. These results are consistent with recent studies showing that unfavorable enzyme-PET interactions can stabilize non-productive states and that appropriately placed interaction motifs can redirect binding toward more productive interfacial configurations [35, 56]. More broadly, they suggest that next-generation PETase engineering should treat interfacial commitment as an explicit optimization target alongside thermostability and catalytic turnover.

The present study also has broader methodological implications. Although Martini 3 has been extensively developed for biomolecular soft-matter systems, its application to protein recognition at synthetic polymer surfaces is still much less systematically benchmarked than its use in proteins, membranes, or small-molecule binding [41–43, 57–61]. Here, the framework reproduces key PET structural and thermophysical properties and supports statistically robust sampling of adsorption and surface-search kinetics over hundreds of independent trajectories. Equally important, it resolves productive configurations in which PET chains enter the catalytic cleft and adopt pre-catalytic geometries. We therefore view this model not simply as a reduced representation of the system, but as a practical mesoscale tool for identifying rate-limiting interfacial events that are difficult to access with fully atomistic simulations at comparable sampling depth.

Several limitations should nevertheless be acknowledged. First, the present simulations resolve adsorption, interfacial search, and pre-catalytic organization, but they do not explicitly model bond cleavage; the PS should therefore be interpreted as a catalytically primed ensemble rather than a chemical transition state. Second, the PET substrate is represented as an amorphous slab with fixed composition, whereas real waste PET contains additional heterogeneity, including crystallinity, surface aging, additives, roughness, and evolving morphology, all of which can influence adsorption and turnover [22,52]. Third, the simulations consider single-enzyme binding events and therefore do not capture collective effects such as surface crowding, competitive adsorption, or coupling to downstream reaction steps. Finally, the designed mutations were evaluated computationally; experimental measurements of adsorption, productive binding, and depolymerization will be needed to quantify how closely the predicted changes in the PS yield translate into improvements in catalytic performance. These limitations affect quantitative transferability, but they do not alter the main conclusion that post-adsorption re-registration and encounter-state over-anchoring are dominant determinants of productive binding.

Overall, our results shift the focus of PETase design from static binding affinity to the kinetic landscape of interfacial commitment. Productive PET recognition emerges not simply from stronger binding, but from a balance between conformational mobility, encounter-state escape, and selective stabilization of docked and pre-catalytic registries. This mechanism-guided perspective provides actionable principles for designing next-generation plastic-degrading enzymes and offers a general framework for understanding enzyme recognition at heterogeneous polymer interfaces.

## 4 Methods

### 4.1 Modeling of PET in Martini 3

#### 4.1.1 CG mapping and bead assignment

The CG representation of PET was constructed following the Martini 3 framework, in which chemically meaningful fragments are preserved and nonbonded interactions are transferred from validated small-molecule building blocks [42, 43]. Each PET repeat unit was decomposed into chemically distinct moieties and mapped to Martini 3 bead types according to fragment size, polarity, and chemical character, while retaining compatibility with Polyply-based polymer construction [45]. Specifically, the ethylene segment (– CH_2_ – CH_2_ –) was mapped to a TC1 bead, and the ester fragment (– C(– O) – O –) was mapped to an SN4a bead. The phenyl ring was represented by two TC5 beads to capture both its size and aromatic character. To account for end-group chemistry, the terminal fragments were treated explicitly: the chain head group – C(– O) – OH was assigned to SP2, whereas the chain tail was represented by N4a for – C(– O) – O – CH_2_ – and TP1 for – CH_2_ – OH. This mapping defines a repeating CG building block suitable for the automated generation of long PET chains.

#### 4.1.2 Bonded interaction parameterization from atomistic reference

Because Martini 3 already provides the nonbonded parameters for the assigned bead types, parameterization focused on the bonded terms, including bonds, constraints, and dihedrals, in order to reproduce the local conformational statistics of PET at the atomistic level. As the atomistic reference, we used a short PET oligomer containing two repeat units (2-PET), which captures the local stereochemistry and torsional preferences governing polymer conformations while remaining computationally tractable.

Atomistic molecular dynamics simulations of 2-PET were performed using the OPLS-AA force field [62], with molecular parameters generated using LigParGen [63–65]. The molecule was solvated in a cubic box of 3.8 × 3.8 × 3.8 nm^3^. After steepest-descent energy minimization, a short *NPT* equilibration run of 0.25 ns was carried out with a 1 fs timestep and all bonds constrained using LINCS. The temperature was maintained at 298.15 K with the velocity-rescale thermostat, and the pressure was maintained at 1.0 bar using the Berendsen barostat. During equilibration, electrostatic interactions were treated using the reaction-field scheme with a 1.4 nm Coulomb cutoff and a reaction-field dielectric constant of 1. Van der Waals interactions were truncated at 1.4 nm, and dispersion corrections were applied to both energy and pressure.

Production simulations were then performed for 500 ns in the *NPT* ensemble with a 2 fs timestep at 298 K and 1.0 bar, using the velocity-rescale thermostat and Parrinello-Rahman barostat. Trajectories were saved every 10 ps. For each snapshot, the atomistic coordinates were mapped to the corresponding CG representation using the mapping scheme in Table S1, and the resulting CG-space distributions of bond lengths, angles (or constrained distances, where applicable), and dihedral angles were extracted.

Bonded functional forms were selected on the basis of both their suitability within the Martini framework and their ability to reproduce the atomistic reference distributions. Bonds and improper dihedrals were described using harmonic potentials, selected stiff degrees of freedom were treated as constraints, and proper dihedrals were represented using the GROMACS Fourier functional form up to fourth order [66]. Starting from initial parameter guesses derived from chemically analogous fragments in the Martini 3 small-molecule database [43], equilibrium values and force constants were iteratively refined to minimize deviations between the atomistic-mapped and CG distributions.

#### 4.1.3 Validation of the CG PET model

The CG PET model was validated at two levels: (i) local conformational statistics and (ii) global solvation/exposure properties.

##### Local conformational statistics

For each bonded degree of freedom, the CG model reproduces the atomistic-mapped distributions with close agreement (Figures S1 and S2). In particular, the calibrated Fourier dihedrals recover the dominant torsional basins required to represent realistic polymer flexibility.

##### Solvent-accessible surface area

As an orthogonal validation metric that was not directly included in the bonded-parameter fitting, we compared the solvent-accessible surface area (SASA) of the atomistic reference and the CG model after mapping. The average SASA differs by only 1.23% between the two representations (Figure S2M), indicating that the CG model preserves the overall exposure characteristics of the oligomer. SASA was computed for both atomistic and CG trajectories using `gmx sasa` [67].

#### 4.1.4 Polyply-compatible force-field definition and PET slab construction

To enable the construction of long PET chains and slab geometries using Polyply, the calibrated 2-PET CG topology was reformatted into a Polyply-compatible Martini 3 polymer definition [45]. Specifically, the PET chain was decomposed into three residue types, `PETinit`, `PET`, and `PETter`, corresponding to the head, repeat, and tail segments, respectively (Table S1). The resulting force-field definition was then added to the Polyply Martini 3 library so that PET could be called as a built-in polymer type.

PET slabs were generated with Polyply in a 10 × 10 × 5 nm^3^ simulation box by first generating the polymer-specific topology with polyply `gen_params` and then constructing the initial coordinates with `polyply gen_coords` using the generated system topology and the target box dimensions [45].

#### 4.1.5 Estimation of the glass transition temperature of bulk PET

To estimate the glass transition temperature (*T*_g_) of the CG PET model, we performed a stepwise cooling protocol adapted from Sangkhawasi *et al*. [68], using the PET slab constructed in the previous step. The system contained 200 PET chains, each comprising 10 repeat units, under periodic boundary conditions.

##### Cooling protocol

Starting from the initial PET slab configuration, the system was cooled from 600 K to 100 K in decrements of 50 K under *NPT* conditions at 1 bar. At 600 K, the structure was first energy-minimized by steepest descent, followed by a 2 ns *NPT* equilibration using the velocity-rescale thermostat and the Berendsen barostat. To implement continuous cooling, each subsequent temperature window was initialized from the equilibrated configuration of the preceding window, including checkpoint-based velocity continuation, and then equilibrated for 2 ns under the same *NPT* conditions. This procedure avoided artificial velocity reinitialization or abrupt temperature rescaling between adjacent windows.

##### Production simulations

After equilibration at each temperature, a 50 ns *NPT* production simulation was performed using the velocity-rescale thermostat and the Parrinello-Rahman barostat. Nonbonded interactions were treated using the Verlet cutoff scheme, with potential-shifted Lennard-Jones and Coulomb interactions truncated at 1.1 nm and a relative dielectric constant of 15.

##### Specific-volume analysis and definition of *T*_g_

At each temperature, the mass density *ρ*(*t*) was extracted from the production energy file. The density was averaged over the final 10 ns of the corresponding 50 ns production trajectory (40-50 ns) to obtain (*ρ*(*T*)), and the standard deviation *σ*_*ρ*_(*T*) was also recorded. The specific volume was then calculated as

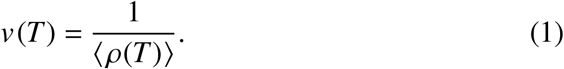

To determine *T*_g_, two independent linear fits were performed for *v*(*T*) in the glassy and rubbery temperature regimes, using the ranges 100-300 K and 400-600 K, respectively. The glass transition temperature was defined as the intersection of these two fitted lines.

### 4.2 vCG PETase simulations

#### 4.2.1 System setup

##### Protein model

The crystal structure of IsPETase (PDB ID: 5XJH) [33] was converted into a Martini 3 CG representation using `martinize2` [69].

##### GōMartini network

To preserve the native fold while allowing large-scale collective motions, a structure-based Gō-like intramolecular network was introduced using the GōMartini scheme. A residue-level contact map was generated from the atomistic reference structure using the Go Contact Map web server [70, 71] and supplied to `martinize2` [69]. Default settings were used for contact-map generation and for assigning the strength of native-contact interactions, consistent with the original GōMartini formulation [40, 44].

##### Complex construction, solvation, and ions

The CG protein was placed above a pre-equilibrated PET slab in a 10 × 10 × 18 nm^3^ simulation box, with the protein centered at (5, 5, 11.5) nm and the slab centered at (5, 5, 2.5) nm (Figure S3). Prior to solvation, the protein was subjected to a rigid-body rotation to avoid strongly back-facing initial configurations, while the initial protein–surface separation remained well beyond the direct interaction range. The system was then solvated with Martini water, and Na^+^ and Cl^−^ ions were added to neutralize the system and achieve a final salt concentration of 0.1 M.

#### 4.2.2 Molecular dynamics simulation protocols

All Martini-based simulations were carried out using GROMACS (version 2024.2) [66] in combination with PLUMED (version 2.9.2) [72, 73] under periodic boundary conditions. Each system was first energy-minimized by steepest descent for up to 20,000 steps, using a convergence criterion of 1000 kJ mol^−1^ nm^−1^ for the maximum force. Production simulations were then performed in the *NPT* ensemble for 1 *µ*s per trajectory using a 5 fs timestep. The temperature was maintained at 298 K using the velocity-rescale thermostat with a coupling constant of 0.5 ps. Separate temperature-coupling groups were used for the protein, solvent/ions, and PET slab. The pressure was maintained at 1 bar using an isotropic Berendsen barostat with a coupling constant of 6.0 ps and a compressibility of 3×10^−4^ bar^−1^. Nonbonded interactions were treated using the Verlet cutoff scheme, with 1.1 nm cutoffs for both Lennard-Jones and electrostatic interactions. Electrostatics were described using the reaction-field method with a relative dielectric constant of 15. Coordinates were saved every 5 ps.

To preserve a single-sided PET-slab geometry along the surface normal, we applied a PLUMED virtual restraint to prevent the protein from reaching the top of the simulation box and interacting with the periodic image of the PET slab. Specifically, an upper wall was imposed on the *z*-component of the displacement between the protein center of mass and a fixed reference point, such that the protein center of mass remained below *z* ≈ 12.5 nm (Figure S3). In practice, the wall was placed at 7 nm relative to the reference point, with a force constant of 1000 kJ mol^−1^ nm^−1^.

#### 4.2.3 Trajectory discretization and state definitions

Each trajectory was discretized into four macrostates (US, ES, DS, and PS) using three scalar observables computed with MDAnalysis [74]: the total protein-PET contact number, CN_total_(*t*); the catalytic-pocket contact number, CN_catalytic_(*t*); and the active-site distance, *D*(*t*). Here, *D* was defined as the Euclidean distance between the side-chain bead (SC1) of the catalytic residue Ser160 and the centroid of the five nearest PET beads. The catalytic pocket was represented by ten residues that define the binding cleft and catalytic region: Tyr87, Trp159, Ser160, Met161, Trp185, Asp206, Ile208, His237, Trp238, and Asn241 (Figure S3B) [33].

Each frame was assigned to one of the four macrostates using the same operational criteria as those described in the main text (Figure 1E). Briefly, the US corresponds to negligible surface contact, with CN_total_ ∈ (0, 1) and the active site remaining distant from the surface, *D* ∈ (1.2, +∞) nm. The ES denotes adsorbed but non-committed configurations, defined by CN_total_ ∈ [1, +∞) and *D* ∈ (1.2, +∞) nm. Productive engagement requires catalytic-pocket contact, with CN_catalytic_ ∈ [1, +∞). Frames satisfying this condition were classified as the DS when *D* ∈ (0.5, 1.2] nm and as the PS when *D* ∈ (0, 0.5] nm.

#### 4.2.4 Visualization and backmapping

All molecular visualization and trajectory rendering were performed using PyMOL (version 0.99) [75] and Visual Molecular Dynamics (VMD, version 1.9.4) [76]. For figures requiring atomistic representations, representative CG snapshots were backmapped to all-atom resolution using CG2AT [77]. Because the PET slab was parameterized here using custom CG bead definitions, an atomistic PET oligomer reference structure and topology were first prepared with the CHARMM-GUI Polymer Builder [78–81]. These files were then used to extend the CG2AT database so that PET could be treated as an integrated polymer during reconstruction. The resulting all-atom structures of the enzyme-PET complex were subsequently exported for publication-quality rendering in PyMOL.

#### 4.2.5 Dimensionality reduction and FEL construction

PCA was performed to characterize the FELs associated with the binding process. For conformational PCA, the analysis was carried out on the protein backbone coordinates after structural alignment to the crystal reference. For contact-space PCA, the input feature vector consisted of the residue-wise contact numbers between surface-exposed protein residues and the PET slab. FELs were reconstructed according to

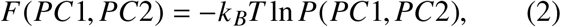

where *P*(*PC*1, *PC*2) is the probability density along the first two principal components.

#### 4.2.6 Mutation protocol for rational design

Point mutations were introduced using the *Mutagenesis* wizard in PyMOL [75]. For each substitution, candidate rotamers were evaluated and the conformation with the fewest steric clashes was selected. The mutated structure was then exported as a PDB file for subsequent CG model construction and simulation.

## Data Availability

The scripts and input files required for setting up the CG MD simulations and performing the subsequent trajectory analyses are publicly available on GitHub at: https://github.com/Gjphysics/PETase_PET_project_code. This repository includes the following components: (1) Simulation Files: GROMACS/PLUMED input files and related setup files for PETase-PET CG simulations. (2) Analysis Tools: Python scripts for trajectory analysis, including contact analysis, state classification, RMSF calculation, PCA, clustering, FEL analysis, and PET slab *T*_*g*_-related calculations.

## Supporting Information

Supporting Information (PDF): Martini 3 mapping scheme for PET and effective trajectory counts; bonded-distance, proper/improper dihedral, and solvent-accessible-surface-area validations of the CG PET model; simulation setup and definition of the IsPETase binding cleft; state-resolved residue-wise protein-PET contact fingerprints and variance-based identification of key residues; state-resolved PCA FELs and RMSF analyses; residue-wise contact fingerprints distinguishing productive and non-productive intermediates; variant-dependent changes in encounter-state contact profiles relative to WT and residue-wise flexibility-contact comparisons; state dwell times and productive micro-pathway kinetics; domain-resolved contact reorganization along the ES→DS reaction coordinate; and mutation-induced changes in state-specific first-arrival times (PDF).

## Acknowledgments

C.H. would like to express deepest gratitude to the Red Bird MPhil Program at the Hong Kong University of Science and Technology (Guangzhou) for providing generous support, resources, and funding, which have been instrumental in the successful completion of this research. X.C. thanks the support from the National Natural Science Foundation of China (Grant No. 12474201), the General Program of the Guangdong Basic and Applied Basic Research Foundation (Grant No. 2024A1515010862), the Guangdong Provincial Project (Grant No. 2023QN10X037) and the Guangdong S&T Program (Grant No. 2025A0505000027). The project was also partly supported by the National Key R&D Program of China (No. 2023YFC3905000). The au-thors acknowledge the Green e Materials (GeM) Laboratory and HPC+AI Intelligence Computing Center at the Hong Kong University of Science and Technology (Guangzhou) for providing computational support.

