## Supplementary Tables and Figures for "Encounter-state over-anchoring governs productive PETase binding on PET surfaces"

### 1 Supplementary Tables

Table S1: **Martini 3 mapping scheme for PET.** Chemical fragments defined for the PET repeat unit and terminal groups, together with the corresponding Martini 3 bead types used in this work for PET slab construction and subsequent contact-based analyses.

| Chemical fragment | Martini 3 bead type |
|---|---|
| $-CH_2-CH_2-$ (ethylene) | TC1 |
| $-C(=O)-O-$ (ester linkage) | SN4a |
| Aromatic ring (phenyl) | 2× TC5 |
| $-C(=O)-OH$ (head end-group) | SP2 |
| $-C(=O)-O-CH_2-$ (tail end-group) | N4a |
| $-CH_2-OH$ (tail end-group) | TP1 |

Table S2: **Numbers of effective trajectories included in the analysis.** $N_{\mathrm{eff}}$ denotes the number of trajectories retained for state assignment, kinetic decomposition, and ensemble-based statistical analyses.

| System | $\mathbf{N}_{\mathrm{eff}}$ |
|---|---|
| WT | 102 |
| ThermoPETase | 103 |
| FAST-PETase | 96 |
| HotPETase | 100 |
| WT-Q119Y | 102 |
| WT-S141F | 100 |
| ThermoPETase-Q119F | 107 |
| ThermoPETase-Y146A | 107 |

#### 2 Supplementary Figures

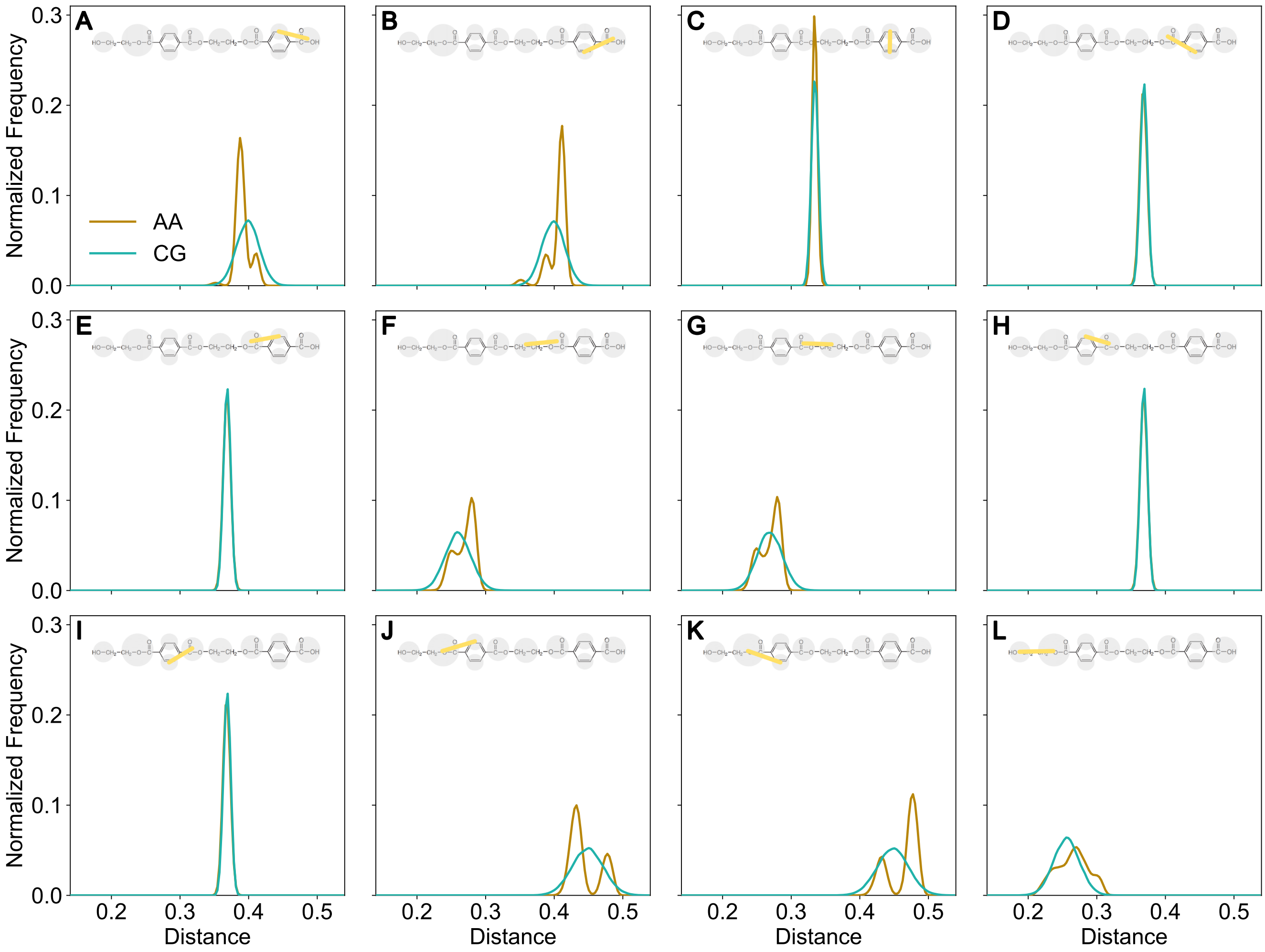


Figure S1: **Calibration of bonded parameters for the Martini 3 PET dimer model.** **(A-L)** Distributions of selected inter-bead distances for the PET dimer (2-mer), comparing the atomistic reference (AA) with the calibrated coarse-grained model (CG). All histograms are shown as normalized frequencies. To balance structural rigidity and conformational flexibility, the AB, FG, and JKL connections were described by harmonic bond potentials, whereas the remaining connections were treated as constraints. The calibrated model reproduces the atomistic reference distributions closely, with constrained distances remaining sharply centered around their target values.

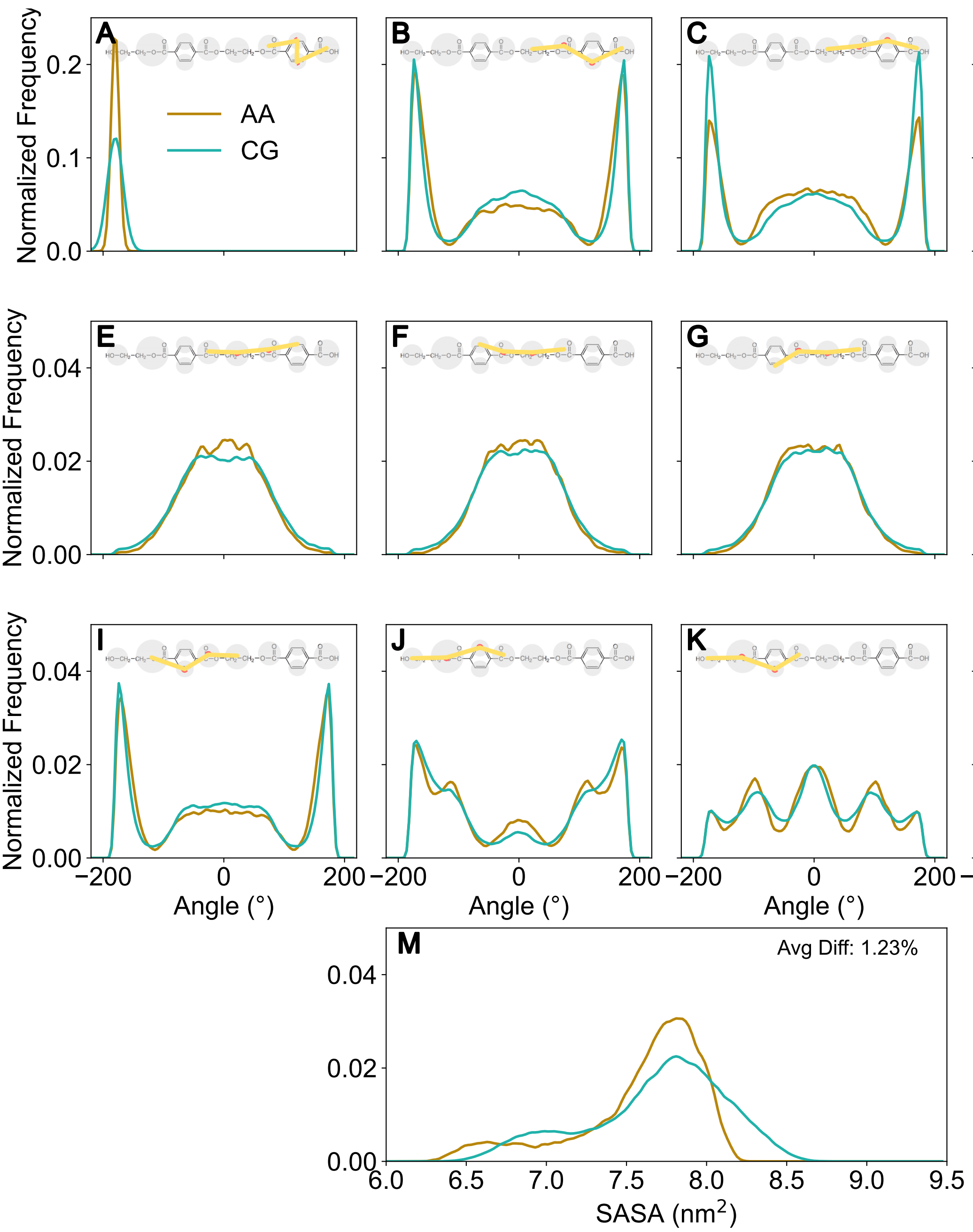


Figure S2: **Calibration of dihedral parameters and SASA validation for the Martini 3 PET dimer model. (B-K)** Distributions of proper dihedral angles for the PET dimer (2-mer), comparing the atomistic reference (AA) with the calibrated coarse-grained model (CG). Proper dihedrals were parameterized using the GROMACS Fourier-type functional form, and the fitted parameters reproduce the major rotameric basins observed in the atomistic reference. **(A, L)** Distributions of improper dihedral angles used to maintain the target local geometry, including the planarity of the corresponding moieties. **(M)** Solvent-accessible surface area (SASA) distributions for the AA and CG models. The average SASA differs by only 1.23% between the two representations, providing an independent validation that the calibrated CG model reproduces the overall conformational ensemble and molecular surface exposure with high fidelity.

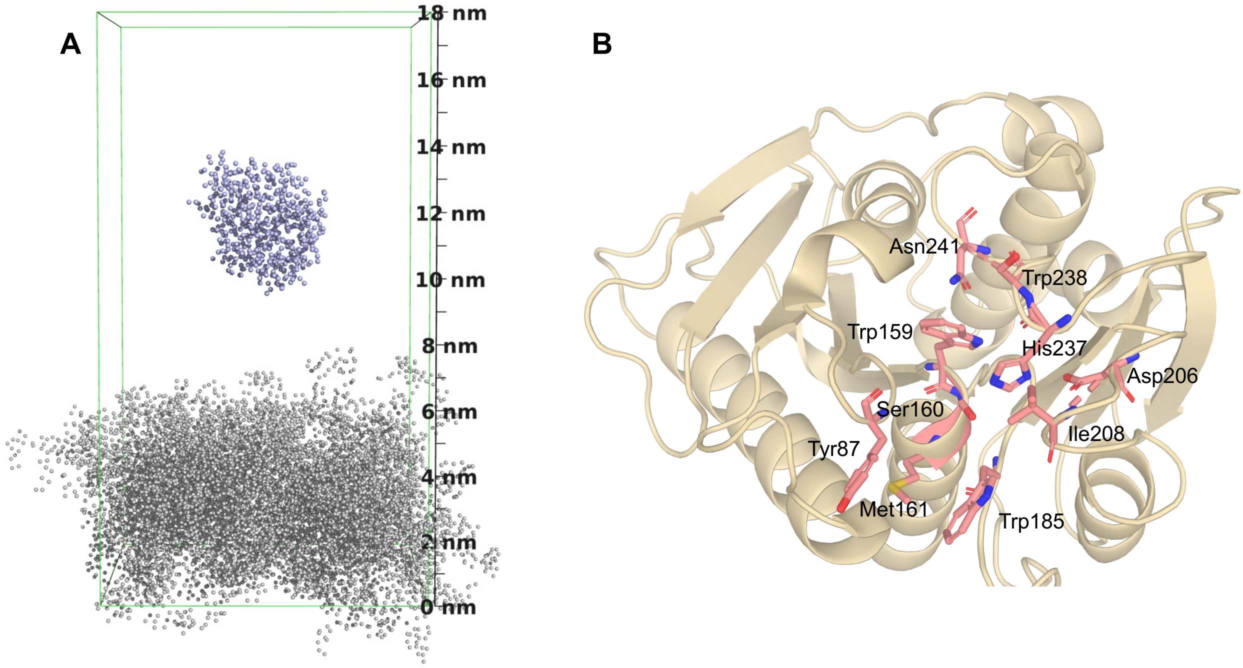


Figure S3: **Simulation setup and definition of the IsPETase binding cleft.** **(A)** Schematic representation of the CG IsPETase-PET slab system. The CG protein was positioned above a pre-equilibrated PET slab in a $10 \times 10 \times 18$ nm$^3$ simulation box, with the protein centered at $(5, 5, 11.5)$ nm and the slab centered at $(5, 5, 2.5)$ nm. To prevent spurious interactions with the periodic image of the PET slab along the $z$ direction, a PLUMED upper-wall restraint was applied so that the protein center of mass (COM) remained below $z \approx 12.5$ nm. **(B)** Residues defining the binding cleft and catalytic region used in the contact and state analyses: Tyr87, Trp159, Ser160, Met161, Trp185, Asp206, Ile208, His237, Trp238, and Asn241.

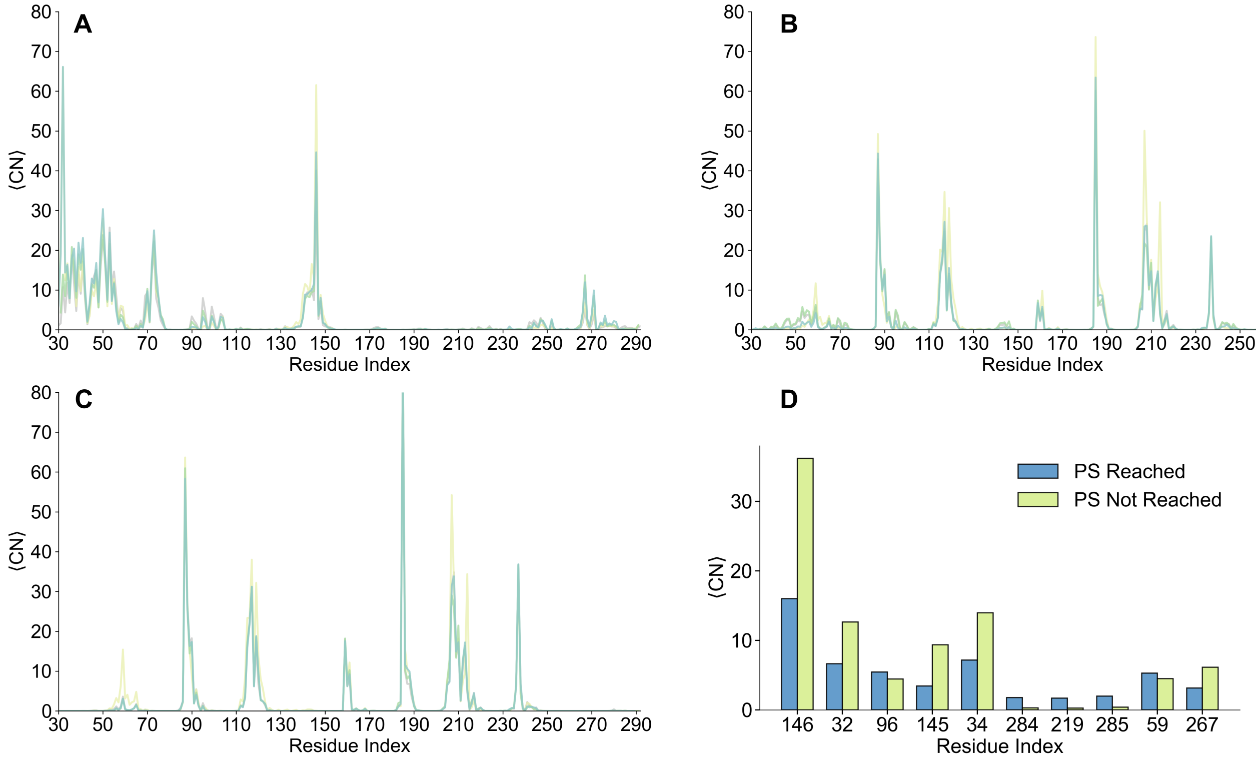


Figure S4: **State-resolved contact fingerprints and variance-based identification of key residues. (A-C)** Ensemble-averaged residue-wise protein-PET contact numbers in the ES, DS, and PS, respectively, shown for WT, ThermoPETase, FAST-PETase, and HotPETase. **(D)** Variance-based ranking of residue importance comparing productive trajectories that reached PS with non-productive trajectories that did not. Trajectories were first divided into PS-reached and PS-not-reached groups according to whether they ever entered PS. For each group, residue-wise protein-PET contact numbers were collected over all frames assigned to ES, and for each residue $i$ an importance score was defined as the absolute difference in contact variance, $I_i = |\mathrm{Var}_{\mathrm{PS}}(cn_i) - \mathrm{Var}_{\mathrm{nonPS}}(cn_i)|$, where $cn_i$ denotes the ES contact number of residue $i$. Residues were then ranked by $I_i$. Bars show the mean ES contact numbers of the top-ranked residues in the two groups, highlighting positions whose contact heterogeneity most strongly distinguishes commitment to the PS.

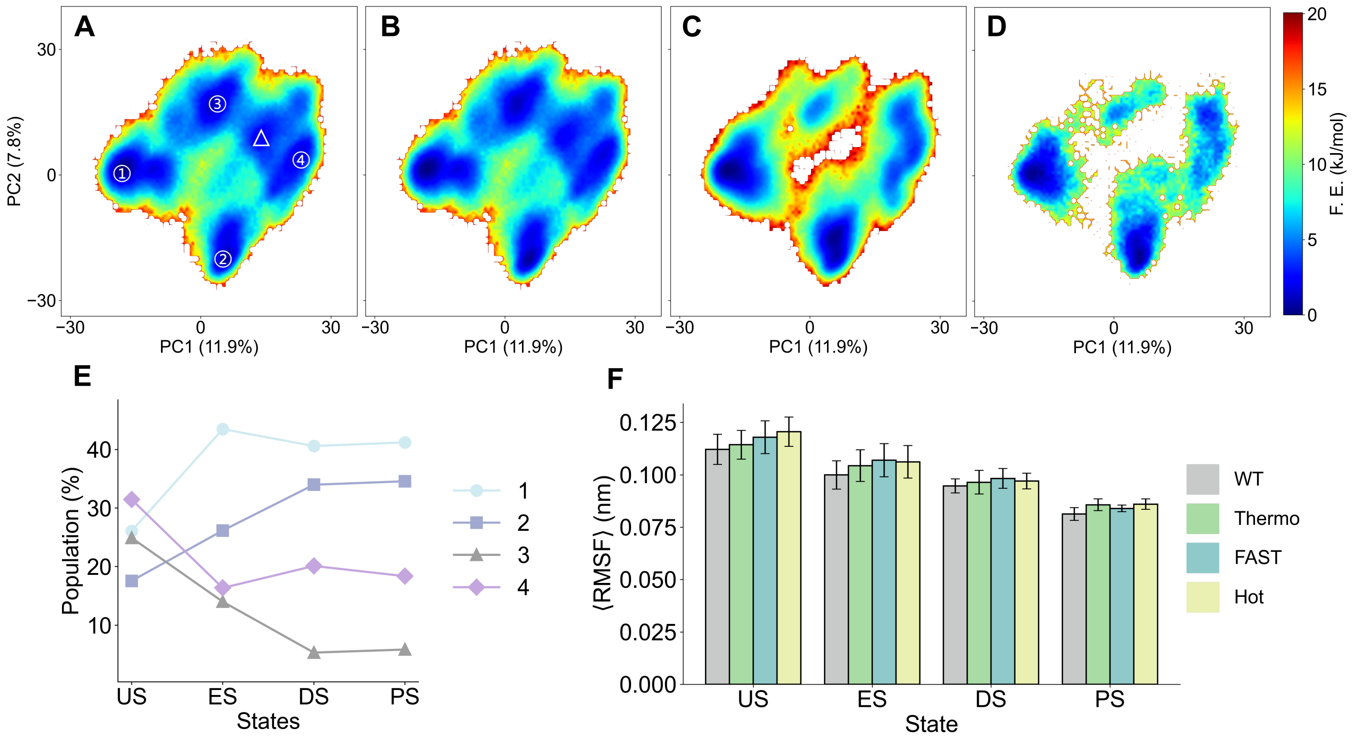


Figure S5: **State-resolved conformational landscapes and flexibility changes along the IsPETase binding pathway.** **(A-D)** Free-energy surfaces of the WT ensemble projected onto the first two principal components (PC1 and PC2), obtained from PCA of all WT trajectories and shown separately for the US, ES, DS, and PS. In **(A)**, labels 1-4 denote the identified local free-energy minima, and the triangle marks the initial configuration used to start the simulations. **(E)** State-dependent populations of the four minima shown in **(A)**, reported as the fraction of frames occupying each basin on the global free-energy landscape. **(F)** Average RMSF values for WT and the engineered variants (ThermoPETase, FAST-PETase, and HotPETase) in different binding states, highlighting the progressive reduction in conformational flexibility during binding maturation.

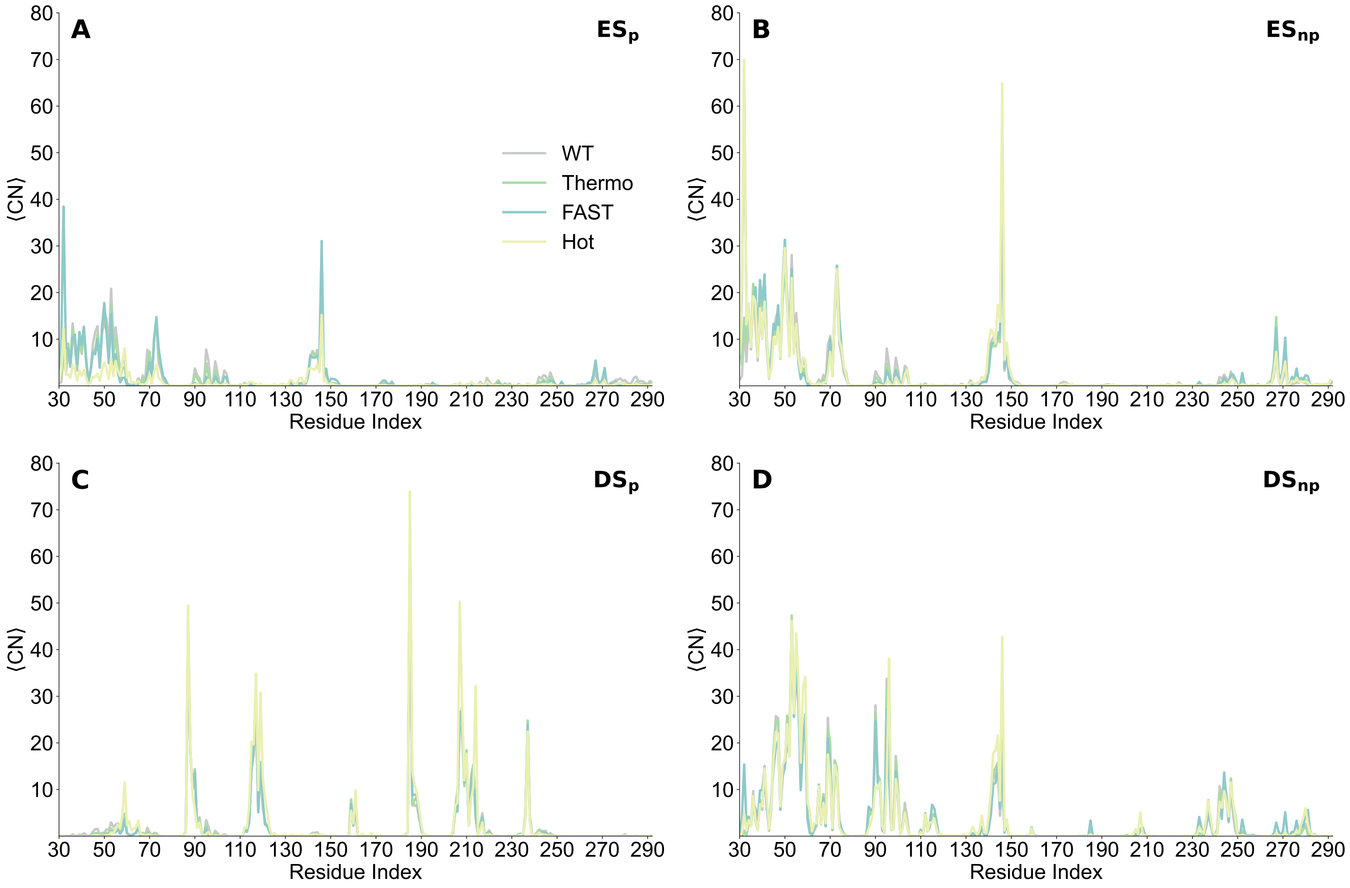


Figure S6: **Residue-wise contact fingerprints distinguishing productive and non-productive binding intermediates.** The average number of protein-surface contacts is shown as a function of residue index for the encounter and docked states. The left column shows productive sub-ensembles that ultimately reach the pre-catalytic state (subscript p), including **(A)** $ES_p$ and **(C)** $DS_p$. The right column shows non-productive sub-ensembles that do not reach PS (subscript np), including **(B)** $ES_{np}$ and **(D)** $DS_{np}$. Differences between the productive and non-productive ensembles identify state-specific residue groups that bias early surface adsorption toward either productive maturation or a competing trapped pathway.

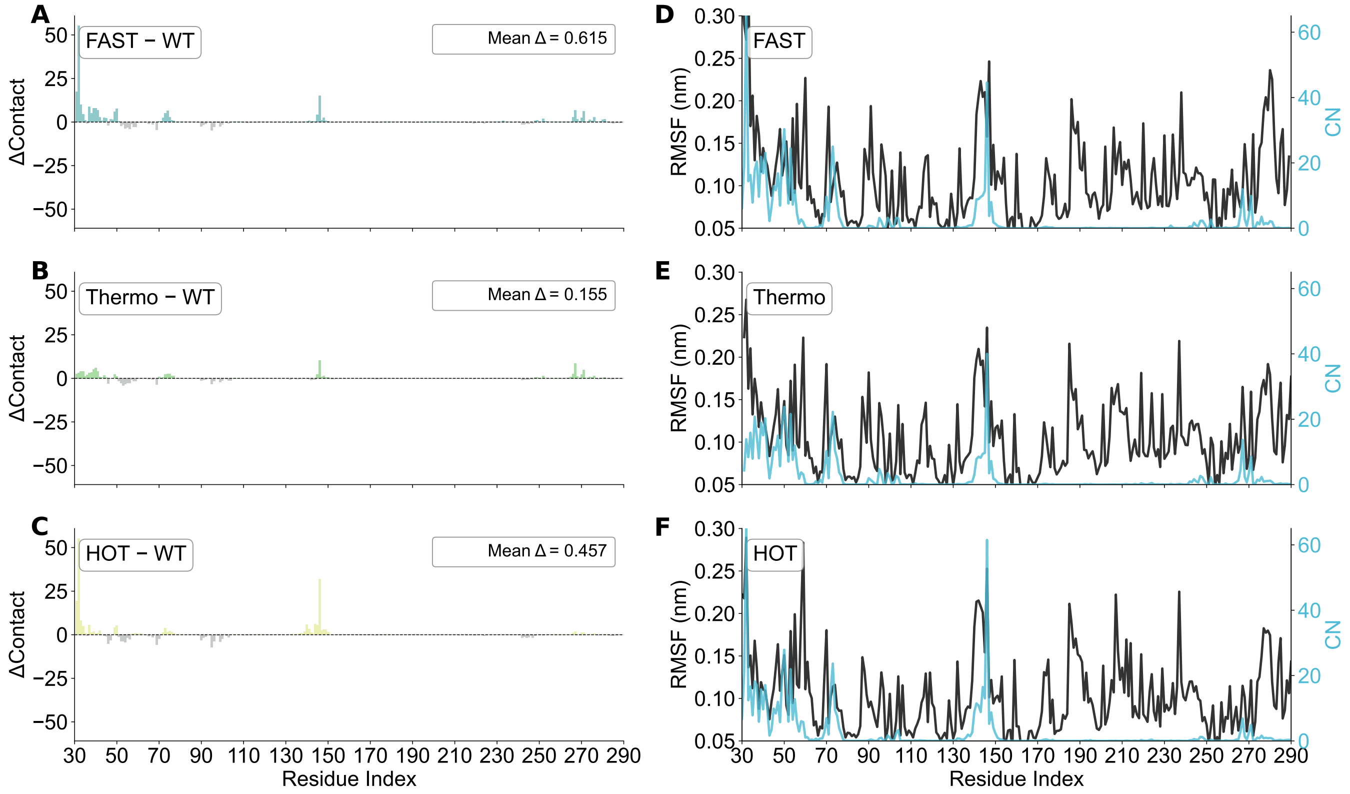


Figure S7: **Variant-dependent changes in ES contact fingerprints relative to WT. (A-C)** Residue-wise differences in the ensemble-averaged protein-PET contact number in the ES, defined as $\Delta CN = \langle CN \rangle_{variant} - \langle CN \rangle_{WT}$, for ThermoPETase **(A)**, FAST-PETase **(B)**, and HotPETase **(C)**. Positive values (colored bars) indicate residues that form more contacts with the PET slab than in WT, whereas negative values (gray bars) indicate reduced contact formation. The mean ES contact shift across all residues is reported in each panel (ThermoPETase-WT: 0.155; FAST-PETase-WT: 0.615; HotPETase-WT: 0.457). **(D-F)** Residue-wise comparison of ES flexibility and PET-contact propensity for FAST-PETase **(D)**, ThermoPETase **(E)**, and HotPETase **(F)**. In each panel, the black curve shows the per-residue RMSF in the ES (left axis), and the cyan curve shows the corresponding ensemble-averaged residue-wise protein-PET contact number (right axis). These comparisons highlight whether strongly contacting regions in ES coincide with relatively flexible or rigid segments.

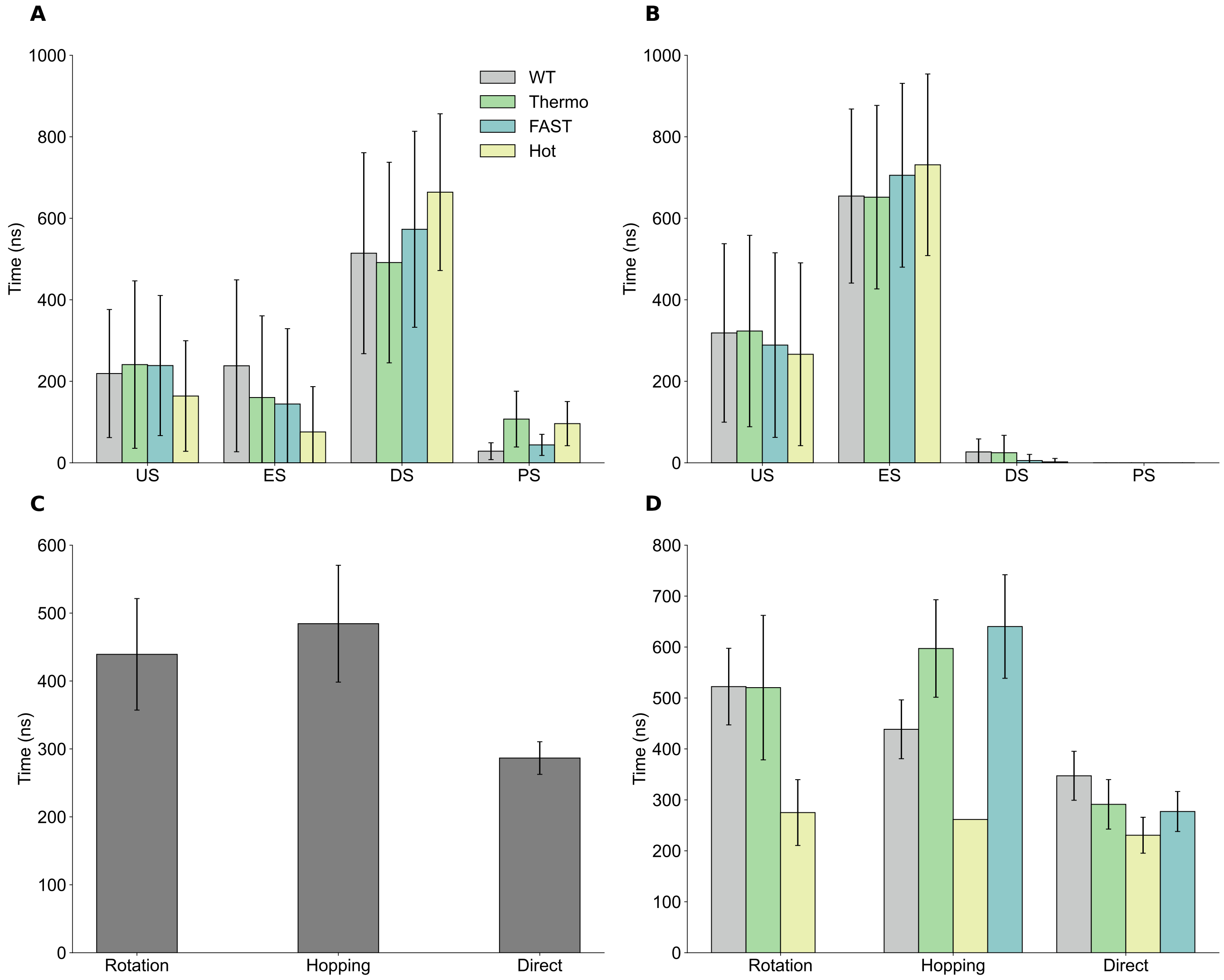


Figure S8: **State dwell times and micro-pathway kinetics for productive and non-productive binding trajectories. (A)** Average time spent in each binding state (US, ES, DS, and PS) for productive trajectories that reach PS, shown for WT and the engineered variants. **(B)** The corresponding state dwell times for non-productive trajectories that do not reach PS within the simulated time window, highlighting prolonged residence in intermediate states and negligible occupancy of PS. **(C)** Average completion times for the three productive micro-pathways, Rotation, Hopping, and Direct, aggregated over all PS-reaching trajectories. **(D)** Micro-pathway completion times resolved by enzyme variant, showing pathway-dependent kinetic differences among WT and the engineered variants. Error bars indicate trajectory-to-trajectory variability and are omitted for groups with insufficient sampling (for example, Hot/Hopping in panel **D**, for which only one trajectory was observed). The Rotation pathway was not observed for FAST-PETase and is therefore not shown.

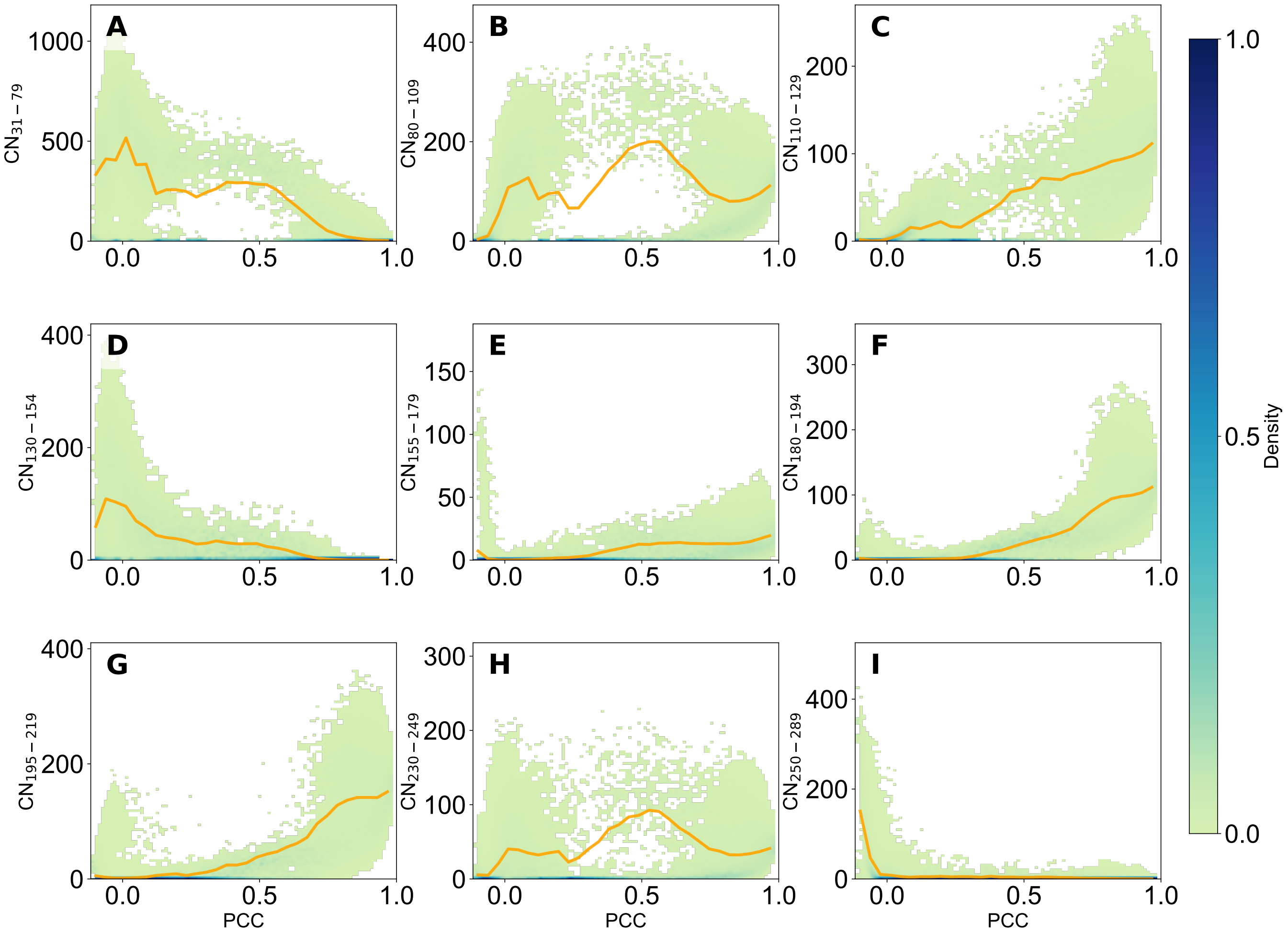


Figure S9: **Domain-resolved contact reorganization along the ES→DS reaction coordinate in WT.** Two-dimensional conditional density plots of the contact sum formed by residues within each domain as a function of the reaction coordinate PCC, defined as the PCC between the instantaneous residue-PET contact profile and the DS reference profile. The WT ensemble is shown for the following residue segments: **(A)** 31-79, **(B)** 80-109, **(C)** 110-129, **(D)** 130-154, **(E)** 155-179, **(F)** 180-194, **(G)** 195-219, **(H)** 230-249, and **(I)** 250-289. The heatmaps represent the conditional distribution of contact sum at a given PCC value, computed from 2D histograms (100 PCC bins × 100 contact-sum bins) after removing NaN/Inf values and excluding frames with near-zero contacts. Histogram counts were normalized independently within each PCC bin, such that the color scale represents $p$(contact sum | PCC) rather than a global density. The orange curve indicates the conditional mean contact sum as a function of PCC.

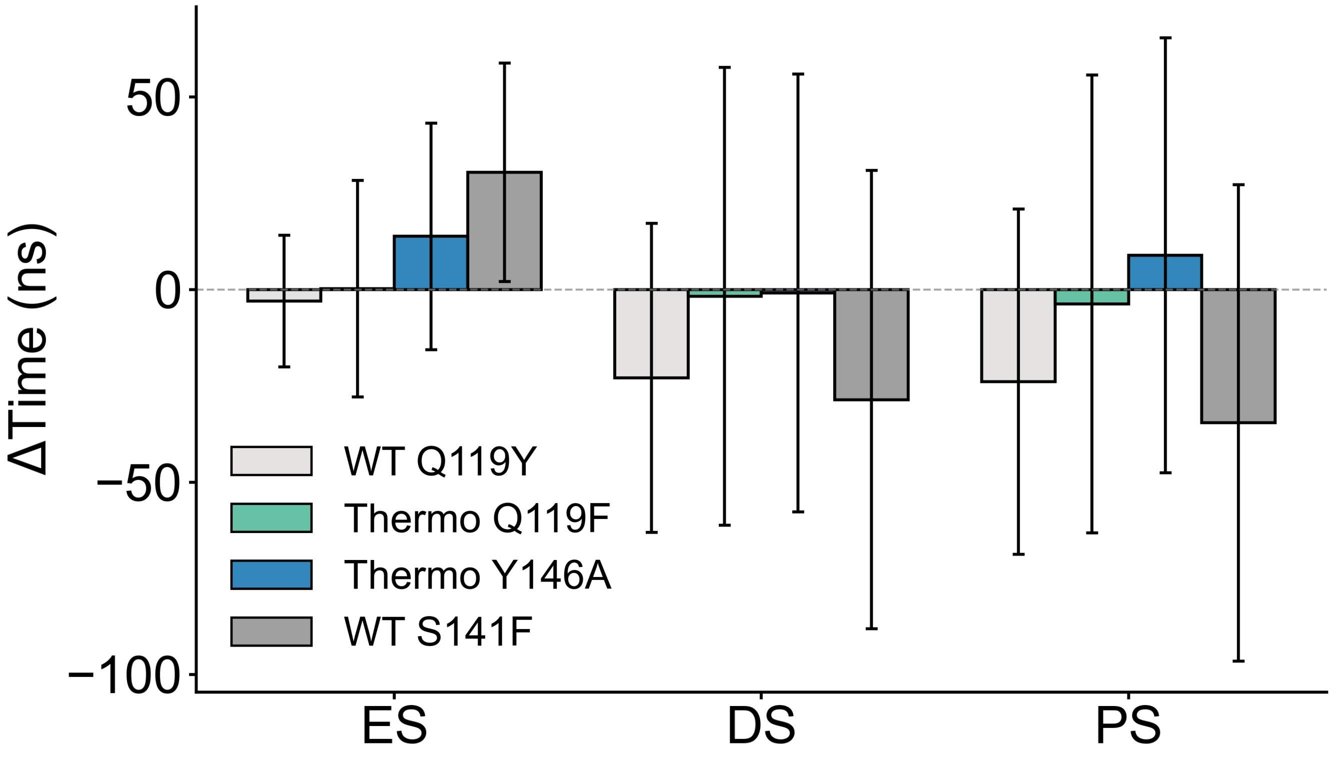


Figure S10: **Effects of rationally designed mutations on state-specific first-arrival times.** Bars show the change in the mean first-arrival time to each binding state (ES, DS, and PS) relative to the corresponding parent scaffold, with the dashed line indicating $\Delta t = 0$. For WT-derived mutants (WT-Q119Y and WT-S141F), $\Delta t = t_{\mathrm{mut}} - t_{\mathrm{WT}}$; for Thermo-derived mutants (Thermo-Q119F and Thermo-Y146A), $\Delta t = t_{\mathrm{mut}} - t_{\mathrm{Thermo}}$. Negative values of $\Delta t$ indicate that the mutant reaches the corresponding state faster than its parent scaffold, whereas positive values indicate delayed arrival.
